# Digital molecular kinetics enabled by large-scale nanopore arrays for multidimensional biophysical profiling

**DOI:** 10.64898/2026.10.28.762428

**Authors:** Kaiju Li, Yan Wang, Yawen Cheng, Tingting Zheng, Shuiying Long, Liping Zhong, Mengdan Wu, Yaozhong Zou

## Abstract

While widely used, traditional technologies like surface plasmon resonance (SPR) are limited by narrow temporal windows (transient capture and baseline drift), dynamic heterogeneity, and poor physiological (37 °C) performance. Meanwhile, early nanopore approaches are often constrained by limited throughput and heavy reliance on customized modification schemes. Here, we present a digital molecular kinetics (DMK) platform based on large-scale nanopore arrays. By introducing a universal coupling and probe reporting system, DMK achieves broad versatility across diverse molecular pairs, including proteins, peptides, small molecules, DNA, and polymerase complexes. In addition to accurately recovering classical SPR association and dissociation rate constants, by harnessing massive single-molecule event analysis, DMK breaks traditional performance boundaries: it facilitates robust thermodynamic-kinetic profiling across a wide temperature range, resolves distinct conformations in heterogeneous samples, enables ultra-low dissociation rate quantification via maximum likelihood estimation, and reveals energy wells via voltage gradients for binding analysis against environmental perturbations. Furthermore, DMK enables the capture of rapid binding events missed by SPR, offering a new perspective on molecular interaction complexity. This platform offers an effective alternative and upgrade to SPR, opening a digital paradigm for complex molecular analysis and drug discovery.

## Introduction

The dynamic interactions of biological macromolecules drive the core mechanisms of life, and quantifying their binding kinetics is essential for fundamental life science, therapeutic development, and more generally, modern biotechnology^1–3^. In scientific history, progress in biological research has largely relied on new analytical tools that push the boundaries of what we can resolve and quantify. As bioscience and its industrial applications rapidly advance, analytical frameworks capable of resolving and quantifying these molecular interactions must advance accordingly to meet increasingly stringent demands for molecular interaction profiling.

For decades, surface plasmon resonance (SPR) has served as the gold standard for biomolecular interaction analysis, playing a pivotal role in the life sciences^4,5^. Nevertheless, as pharmaceutical and biotechnological research advances to demand multidimensional and high-resolution thermodynamic and kinetic profiling, its optical-fluidic platform and population-level averaging encounter intrinsic technical limitations. Constrained by thermal drift, direct acquisition of physiological-temperature kinetic profiling is difficult, and reliance on extrapolations often distorts accuracy^4^. Additionally, its macroscopic optical signal obscures conformational heterogeneity and rapid transient states^6,7^. In kinetic quantification, mass-transport and system response limits can constrain the resolution of ultrafast association and dissociation, whereas baseline drift complicates ultra-slow dissociation measurements^8^. Finally, its laminar flow configuration is restricted to near-zero-force regimes, leaving open the need for probing mechanical energy landscapes and physiological shear responses.

Single-molecule technologies have garnered widespread attention for mitigating ensemble-averaging and resolving conformational heterogeneity. Among various platforms, nanopores have emerged as powerful tools for capturing transient molecular interactions by recording ionic current signals with high temporal resolution. Traditional nanopore interaction assays fall into three classical strategies: volume-exclusion pore blocking, where complex formation occludes the pore and generates characteristic current interruptions^9–11^; pore-interior immobilized capture, which traps analytes inside the pore via specific modification and changes the current signal accordingly^12–14;^ and spatially confined recognition, where the target molecule is conjugated in the confined sensing area of the nanopore, which relies on binding-induced conformational changes within a restricted sensing zone to generate specific signals^15–17^. All three approaches heavily rely on the charge properties, size, and geometrical adaptation of molecules to the pore structure, limiting their generalizability. Thakur and Movileanu (2018) introduced a probe-assisted strategy to decouple signal generation from the direct properties of target molecules^18^. By anchoring receptor-probe fusions near the pore entrance, this approach transduces binding events through target-induced conformational changes that modulate probe-nanopore interactions. However, this mechanism remains constrained by the required probe conformational changes, forcing custom-screening of fusion sites for every new target. The tedious engineering of this "one pore, one target, one probe" paradigm, combined with the throughput limitations of traditional single-channel platforms, fundamentally prevents it from supporting large-scale, statistically robust universal analysis.

Here, we present a single-molecule interaction platform combining broad versatility with high throughput. In addition to conventional fusion expression, it constructs functional nanopore-target and probe complexes via standardized chemical conjugation, eliminating the "one pore, one target, one probe" engineering paradigm. Utilizing commercial high-density parallel nanopore arrays, the platform simultaneously captures tens of thousands of single-molecule events. In contrast to traditional SPR relying on macroscopic optical signals, absolute digital counting of discrete single-molecule interactions resolves molecular and kinetic heterogeneities, while high time-resolution and baseline stability extend the kinetic window to enable precise quantification from ultra-fast binding to ultra-long residence times. Supported by real-time temperature control and electric field regulation, it resolves physiological-temperature kinetics and mechanical energy landscapes. By integrating multi-dimensional profiling, broad versatility, and scalability, this platform provides a powerful framework to advance biomolecular interaction analysis.

## Results

### 1. Detection Mechanism

Fig. 1 illustrates the design of our platform utilizing nanopores as signal sensors for molecular interaction kinetics. To eliminate the confusion and ambiguity arising from context-dependent biochemical nomenclature, we uniformly define the interacting pair into two designated entities: Partner A (anchored to the nanopore via a tether) and Partner B (free in solution). To ensure system versatility, the coupling between Partner A and the nanopore can be achieved through standardized chemical conjugation as well as conventional fusion expression, allowing Partner A to be readily anchored to nanopores pre-functionalized with a reactive handle at a specific site. The probe molecule consists of a charged polypeptide, and this uniform architecture can be conjugated to various Partner B molecules via click chemistry to maximize platform generalizability. Upon the binding of Partner B to Partner A, the probe is brought into close proximity to the nanopore entrance (typically within ∼10 nm), creating a millimolar-scale local effective concentration that enables efficient capture of the probe by the nanopore under an electric field, generating characteristic current blockades that serve as the foundation for signal detection. In contrast, free Partner B molecules, due to their much lower bulk working concentrations (typically well below the micromolar level, down to the low nanomolar range), exhibit capture efficiencies that are several orders of magnitude lower. With high-frequency alternating current (AC) stimulation, probe molecules in the bound state are repeatedly captured throughout the binding duration to produce sustained characteristic signals. Conversely, free probes, even if occasionally captured, escape from the nanopore and diffuse away under the reverse voltage, generating only sporadic interference that is readily filtered out by standard signal processing algorithms.

**Fig 1.**
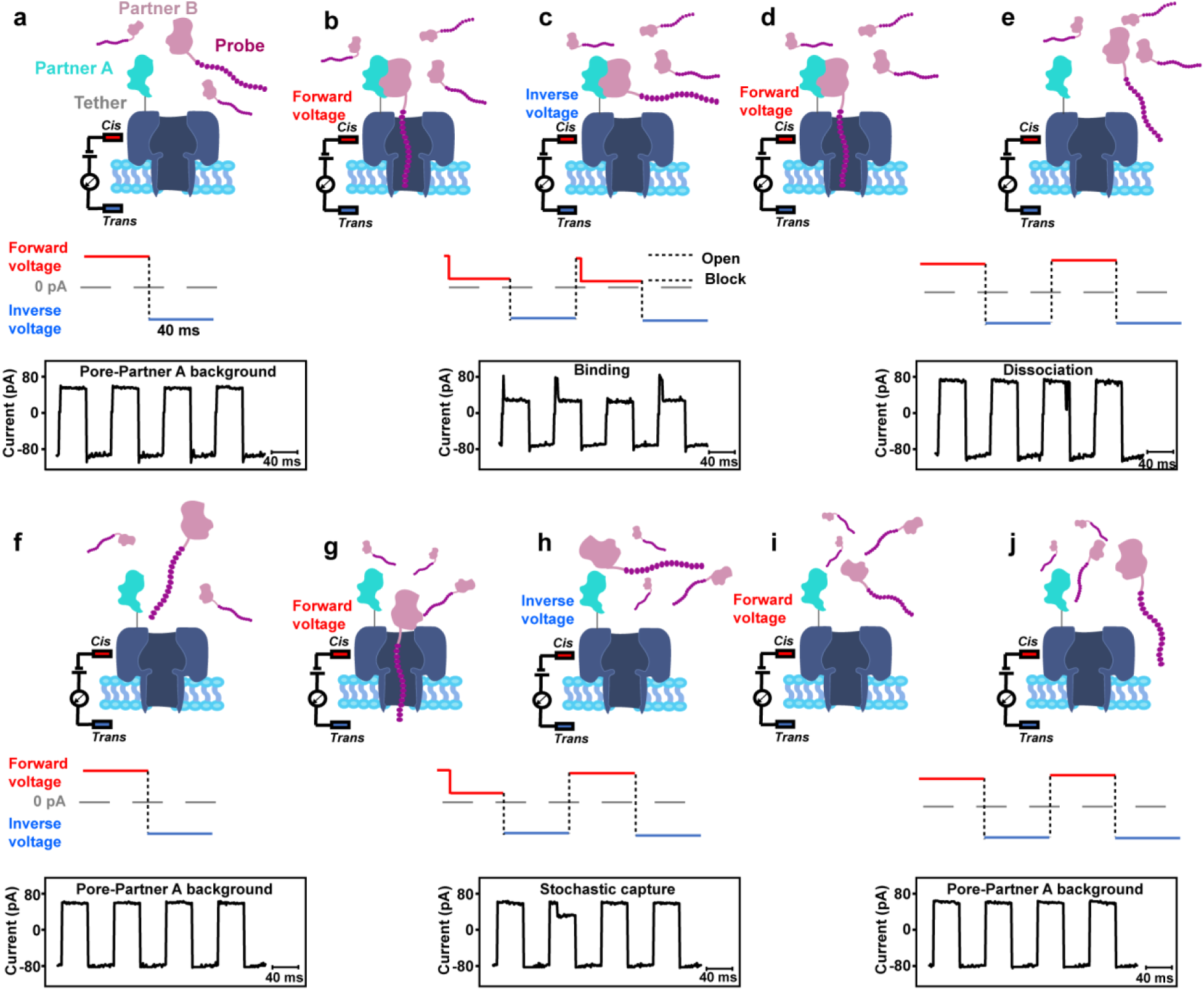
Principle of molecular interaction sensing via nanopore-probe technology. **a–e,** Characteristic cross-cycle blockade signals generated upon Partner B binding to Partner A: **a,** Partner A is tethered to a monomer of the nanopore via a linker to form a Pore–Partner A complex; a charged peptide probe is conjugated to Partner B in solution. The complex exhibits an open-pore background current under AC voltage excitation. **b,** Upon binding of Partner B to Partner A, the probe attached to Partner B is captured by the nanopore under forward voltage, producing a specific current blockage. **c,** Under reverse voltage, the probe is ejected from the nanopore (current returns to the open state) while the Partner A–Partner B complex remains intact. **d,** Under forward voltage in the subsequent cycle, the probe is recaptured into the nanopore, yielding repetitive, cross-cycle current blockades. **e,** Upon dissociation, the current restores to the open state. **f–j,** Stochastic capture events of uncomplexed B-probe: **f,** Same as in a, open-pore background current. **g,** Under forward voltage, the probe may be captured via stochastic collisions, producing a characteristic blockage event. **h,** Under reverse voltage, both the probe and Partner B are ejected and diffuse away. **i,** Due to low local concentration and capture probability, the probe is no longer captured in the next cycle, returning to the open-pore state. **j,** Uncaptured background state, same as in a and **f.**

Compared with traditional nanopore sensing, this scheme features three distinct advantages: First, modular and standardized coupling schemes significantly enhance system versatility, overcoming the constraints of conventional "one pore, one target, one probe" formats. Second, the core signal mechanism requires neither conformational changes nor specific modification sites; instead, it relies on the binding-induced elevation in the local effective concentration of the probe relative to the nanopore, conferring unparalleled generalizability. Third, high-frequency AC effectively filters free probe signals and enables reliable discrimination of true binding events from background interference. In addition, it circumvents the pore-clogging drawbacks inherent to traditional nanopore detection under direct current (DC) stimulation.

### 2. Ensemble Kinetics and Digital Single-Molecule Detection

The distance-dependent sensing mechanism can also be integrated with large-scale nanopore arrays on microfluidic chips for high-throughput parallel detection. Implemented on the G-seq500 device equipped with the MK02 chip (featuring 512K addressable channels)^19^, this platform readily supports the robust integration of Partner A-conjugated nanopores and the subsequent automated analysis of Partner B binding, enabling massively parallel electrical readout (Fig. 2a). In this setting, the tens of thousands of Partner A-conjugated nanopores serve as analogues of ligands immobilized on a SPR sensor chip, while the collective electrical blockade signals correspond to the optical refractive index responses in SPR. By monitoring the fraction of nanopores exhibiting elevated blockade states (termed blocking rate) during the introduction of Partner B and subsequent buffer washout, we readily generate real-time kinetic curves similar to SPR sensorgrams (Fig. 2b). With standard SPR-compatible data analysis workflows, quantitative kinetic parameters can be reliably extracted, yielding kinetic results directly comparable to traditional SPR data.

**Fig 2.**
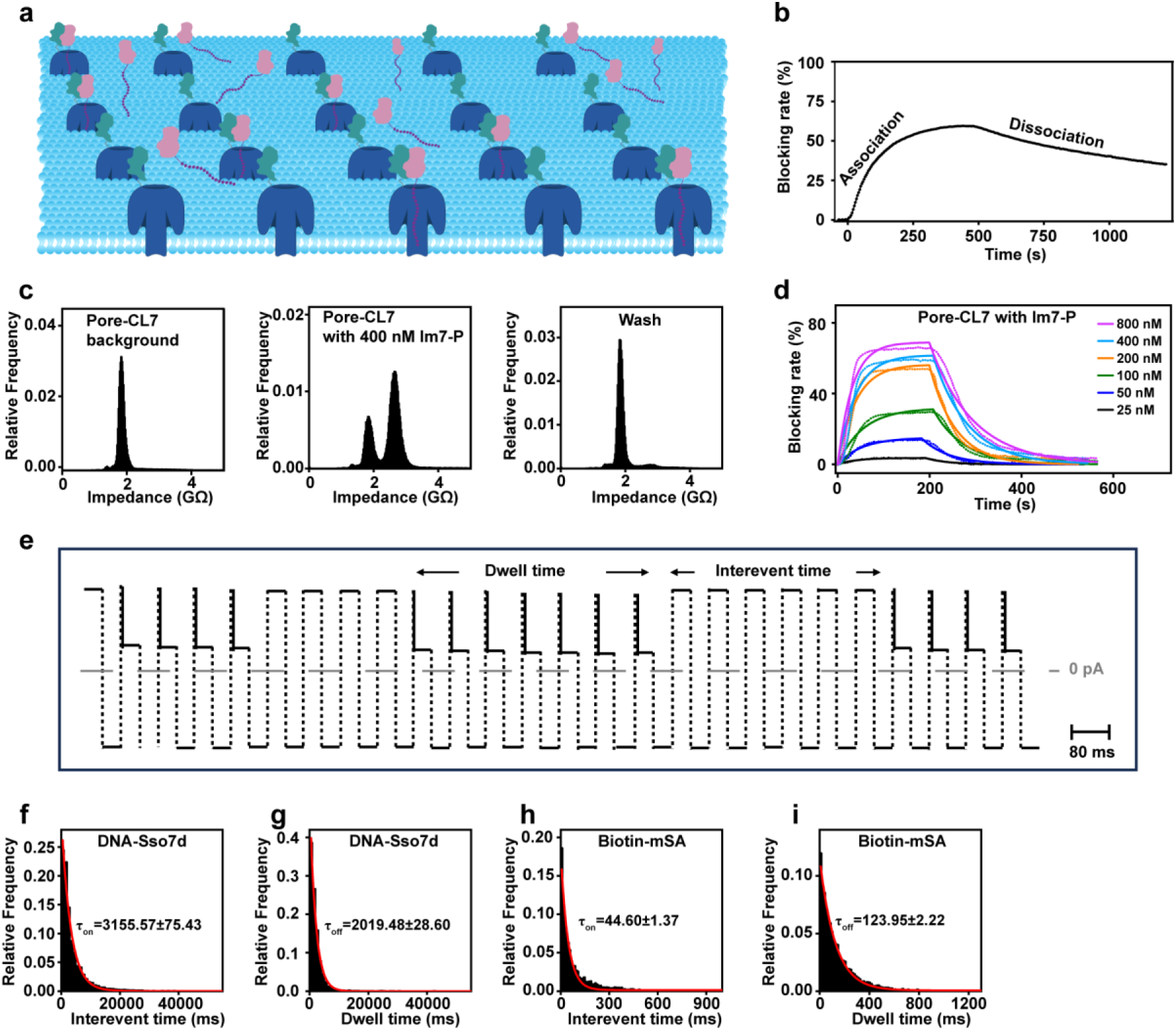
Quantifying molecular interaction kinetics using high-throughput nanopore arrays. **a,** Schematic illustration of the nanopore-probe sensing mechanism (Fig. 1) integrated into a high-throughput array format. **b,** Ensemble binding-dissociation curves derived from real-time changes in blocking rate over time, analogous to macroscopic SPR sensorgrams. **c,** Impedance distribution histograms of the Pore-CL7 complex interacting with Im7-P. Left: Single open-pore low-impedance peak for Pore-CL7 alone. Middle: Appearance of a high-impedance blockage peak upon addition of 400 nM Im7-P to the cis side. Right: Upon washing, the fraction of the high-impedance blockage peak is markedly reduced due to analyte dissociation and clearance, accompanied by the restoration of the open-pore peak. **d,** Real-time ensemble binding-dissociation curves at indicated Im7-P concentrations recorded across the array, fitted with a 1:1 binding model. The blocking rate is derived from the real-time fraction of blocked relative to total active pores. **e,** Representative trace showing the extraction of single-molecule interaction events. The duration of each blockage event (dwell time) reflects the bound-state lifetime for *K_off_* determination, whereas the time interval between consecutive events (interevent time) corresponds to the binding frequency for *K_on_* determination. **f, g,** Statistical distributions and single-exponential fits of interevent times **(f)** and dwell times **(g)** for the interaction between DNA and Sso7d-P (at 40.8 nM), yielding the characteristic times *τ_on_* and *τ_off_*. **h, i**, Statistical distributions and single-exponential fits of interevent times **(h)** and dwell times **(i)** for the interaction between biotin and mSA-P (at 6.25 nM), yielding the characteristic times *τ_on_* and *τ_off_*. All experiments were performed in 200 mM KCl, 50 mM phosphate buffer (pH 6.8) under an alternating voltage of ±100 mV.

We first tested the protein-protein interaction between colicin E7 DNase (CL7) and its cognate immunity protein (Im7)^20,21^. CL7-conjugated nanopores were incorporated into the chip (see Methods for details) and initially exhibited a uniform open-pore state with an impedance of approximately 1.85 GΩ (Fig. 2c left). Upon the addition of the Im7-probe, specific binding to CL7 facilitated the capture of the probes, yielding an elevated blockade state of approximately 2.65 GΩ (Fig. 2c middle). The fraction of these high-impedance channels displayed clear concentration dependence as well as exponential time dependence, intuitively tracking the dynamic association process of CL7-Im7. Similarly, during the subsequent buffer washout, the fraction of high-impedance channels exhibited an exponential decay profile characteristic of dissociation kinetics (Fig. 2c right), yielding macroscopic kinetic trajectories closely resembling SPR sensorgrams (Fig. 2d). Fitting these curves to a 1:1 binding model enabled determination of kinetic parameters for the CL7-Im7 interaction (Supplementary Table 1). These values are in close agreement with classical SPR results, demonstrating the validity of our platform as a viable alternative to SPR (Supplementary Fig. 1 and Supplementary Table 2).

Employing a similar approach, we further determined the kinetic parameters for several additional model pairs, including an affinity-enhanced mutant CL7 (CL7_M)^22^ and Im7, monomeric streptavidin (mSA) and biotin^23^, Single Stranded DNA Binding Domain 7D (Sso7d) and DNA^24^, and SpyTag and SpyCatcher^25^. The results are all in close agreement with those obtained by SPR or reported in the literature (Supplementary Fig. 2 and Supplementary Table 1-2). As the transient Sso7d-DNA interaction with rapid kinetics cannot be resolved by conventional SPR-like analysis, only the equilibrium dissociation constant (*K_d_*) was determined (Supplementary Fig. 2 e-g). Conversely, the binding of SpyTag and SpyCatcher involves the formation of a covalent isopeptide bond^26^, our system enables the quantitative characterization of its covalent coupling kinetics (Supplementary Fig. 2 c-d). For the mSA-biotin pair, we tested reciprocal conjugation strategies—alternating which component was conjugated to the nanopore versus labeled with the probe. Both strategies were successful and generated highly consistent kinetic parameters (Supplementary Fig. 2h-k and Supplementary Table 1). The accurate characterization of these diverse biomolecular interactions comprehensively demonstrates the broad versatility of our platform.

Single-molecule detection in real-time via nanopore arrays can track individual association and dissociation events across tens of thousands of molecular pairs in parallel. As shown in Fig. 2e, each sustained blockade signal corresponds to the duration of a binding event (dwell time), which can be used to fit the dissociation rate constant (*K_off_*). Conversely, the interevent time between successive binding events dictates the association kinetics, enabling the fitting of the association rate constant (*K_on_*). Moving beyond traditional macroscopic signal intensities, this strategy of converting single-molecule association and dissociation events into digital electrical signals with high temporal resolution inspires the designation of our platform: Digital Molecular Kinetics (DMK).

For rapid, transient binding systems such as Sso7d and DNA, traditional macroscopic kinetic analysis methods often struggle to resolve kinetic constants and only *K_d_* can be determined; however, enabled by the precise single-molecule tracking and fitting capabilities of DMK (Fig. 2 f-g), we successfully extracted the *K_on_* and *K_off_* values that remain elusive to conventional methods (Supplementary Table 3). The *K_d_* values derived from DMK kinetic parameters show close agreement with results from macroscopic ensemble kinetic analyses, demonstrating the internal consistency of our platform bridging macroscopic and single-molecule regimes. Beyond the Sso7d-DNA system, we further applied the DMK strategy to determine the kinetics for mSA and biotin (Fig. 2h-i and Supplementary Table 3). Compared with macroscopic kinetic results, the *K_d_* values exhibit high consistency, whereas minor, fold-level differences are observed for *K_on_* and *K_off_* (displaying slightly faster absolute values while remaining within the same order of magnitude). This deviation can be ascribed to experimental error and possible local rebinding effects induced by inevitable mass transport limitations and imperfect perfusion efficiency in macroscopic fluidic systems, particularly for high-*K_on_* interactions. Furthermore, macroscopic kinetic methods may miss transient binding events of extremely short duration due to limited response bandwidth, which could also partly contribute to the underestimation of macroscopic *K_on_* and *K_off_* values.

### 3. Thermodynamic Parameters and Physiological-Temperature Kinetics

Thermodynamic parameters including changes in Gibbs free energy (Δ*G*), enthalpy (Δ*H*), and entropy (Δ*S*) elucidate the underlying driving forces of molecular interactions. Isothermal titration calorimetry (ITC) is widely recognized as the gold standard for determining binding thermodynamics; however, it is limited by expensive instrumentation, high sample consumption, and low signal-to-noise ratios when assessing weak thermal reactions^27^. Theoretically, based on the van’t Hoff equation (Equation 1), thermodynamic parameters can be derived from kinetic constants obtained across various temperatures^28,29^. However, SPR is acutely sensitive to temperature fluctuations and severe thermal drift during variable-temperature operations renders the reliable acquisition of multi-temperature kinetic data exceptionally difficult in practice.

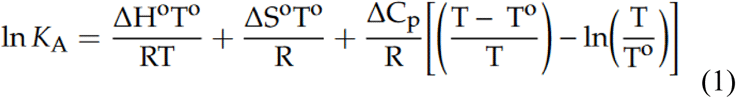

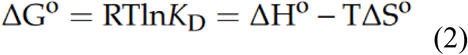

By contrast, the hardware of our system based on nanopore sensing achieves precise temperature control within the 15-40 °C range on a timescale of seconds, removing technical barriers to variable-temperature experiments. Taking the CL7-Im7 system as an example, we determined the kinetic constants across the 13-37 °C range using the DMK strategy (Supplementary Fig. 3 and subsequently derived the corresponding association constants *K_A_* (Fig. 3a). Based on van’t Hoff analysis and the Gibbs free energy equation (Equation 2), fitting parameters by quadratic polynomial (Supplementary Fig. 4, we successfully obtained the complete set of thermodynamic parameters for CL7-Im7 (Fig. 3c). The Δ*G* shows near-perfect agreement with that from ITC (Fig. 3c and Supplementary Fig. 5), and similar Δ*H* and Δ*S* values indicate that CL7-Im7 binding is a typical enthalpy-driven process accompanied by a significant entropy penalty, demonstrating the unique value of the DMK platform in resolving molecular thermodynamic mechanisms.

**Fig 3.**
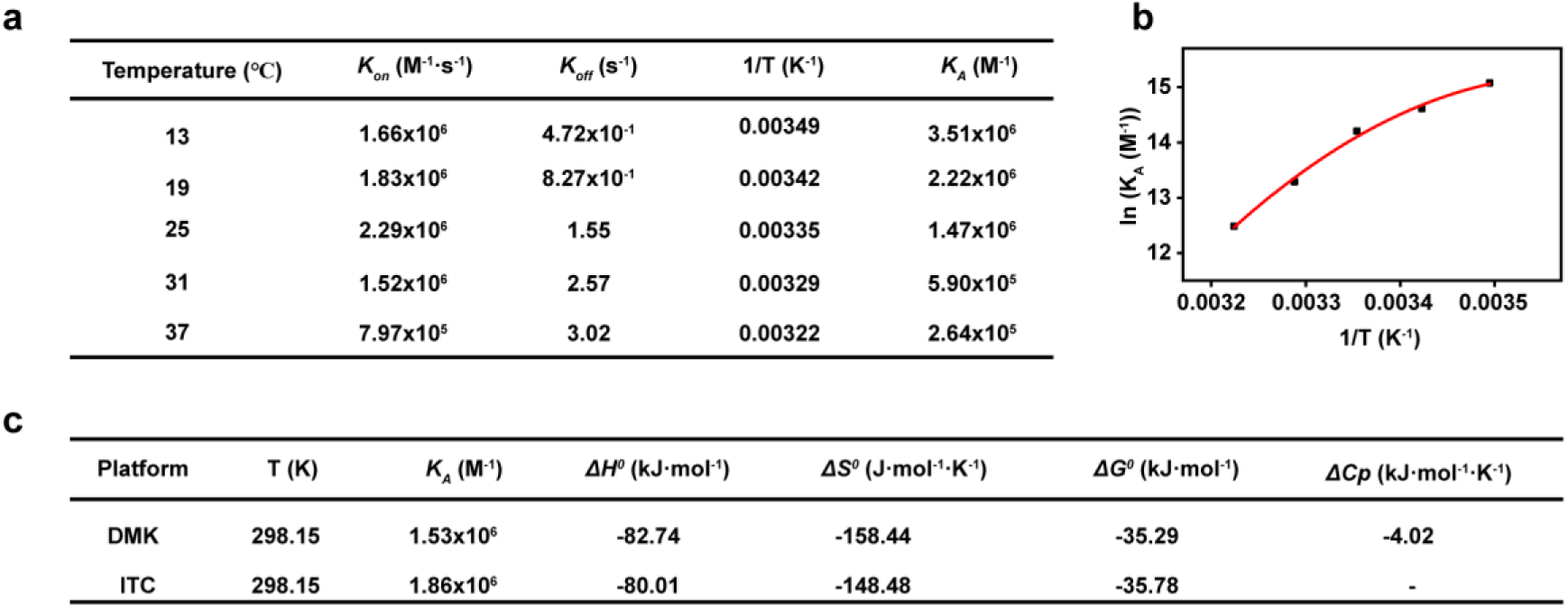
Thermodynamic characterization using the DMK platform. **a,** Kinetics of the CL7-Im7 interaction measured by DMK at various temperatures (13-37 °C). **b,** Van’t Hoff plot showing the equilibrium association constant (*K_A_*) as a function of reciprocal temperature (1/T) derived from DMK kinetics at different temperatures. Data were fitted to a second-order polynomial equation to determine Δ*H*, Δ*S*, Δ*G*, and Δ*Cp* values according to Equations (1) and (2). **c,** Comparison of thermodynamic parameters for the CL7-Im7 interaction obtained from DMK and ITC. These experiments were performed in 200 mM KCl, 50 mM phosphate buffer (pH 6.8) at a voltage of ±100 mV.

Notably, as temperature increases, the *K_off_* and *K_A_* values for CL7-Im7 evolve rapidly, exhibiting pronounced non-linearity within the 13-37 °C range (Fig. 3b). Consequently, van’t Hoff fitting forces an anomalously large heat capacity change (Δ*Cp*) term. However, this inflated value is physically implausible as a genuine heat capacity alteration; instead, it more likely reflects temperature-induced conformational adjustments, as supported by kinetic reversibility upon temperature resetting (Supplementary Table 4). This limitation of van’t Hoff extrapolation partly accounts for the slight deviations between our derived Δ*H* and Δ*S* values and the ITC benchmark. While such thermodynamic deviations remain acceptable for qualitative profiling, attempting to extrapolate kinetic behavior to physiological temperatures may introduce substantial error due to this pronounced non-linearity, underscoring the necessity of direct measurement at 37 °C for pharmacological and industrial applications.

### 4. Kinetic Analysis of Heterogeneous Samples

Sample heterogeneity, such as distinct conformational states or post-translational modifications, poses a constant challenge to macroscopic platforms like SPR and ITC. Ensemble averaging inherently obscures these molecular complexities, rendering them unable to resolve the distinct dynamic behaviors of subpopulations that may modulate physiological or pharmacological outcomes. Based on single-molecule sensing, the DMK system possesses a fundamental advantage in resolving heterogeneous samples^9,30^. Taking the interaction between Bim and its binding peptide (Bim binder) as an example^31^, SDS-PAGE reveals two distinct bands for the Bim binder (Supplementary Fig. 6) and functional validation confirms that both bands retain binding activity toward Bim (Fig. 4a), suggesting that they correspond to distinct functional subpopulations. Correspondingly, in DMK kinetic assays, nanopore channels functionalized with Bim binder exhibit two distinct residence time distribution patterns (Fig. 4b and Methods). Partitioning these channels accordingly allows us to obtain two distinct sets of kinetic parameters (Fig. 4c-h and Supplementary Table 6), yielding a subpopulation ratio (73.43% : 26.57%) that shows high consistency with the densitometric quantification of SDS-PAGE bands (78.49% : 21.51%, Supplementary Table 5). This result demonstrates the capability and reliability of the DMK system in resolving kinetic parameters for heterogeneous samples. In contrast, traditional SPR was mathematically ill-suited to resolve the heterogeneity of the Bim binder, yielding either a homogenized single set of kinetic constants or physically implausible kinetic parameters when forced into multi-state models (Supplementary Fig. 7 and Supplementary Table 7). Notably, this averaged constant derived by SPR aligns closely with the kinetics of the slow-dissociating subpopulation resolved by DMK (Supplementary Table 6 and 7), providing additional corroboration for the reliability of the DMK-derived parameters. Furthermore, even if SPR could successfully fit the data, it remains fundamentally incapable of validating the biophysical reality of multi-exponential or heterogeneous models. By comparison, the DMK system achieves in situ resolution of heterogeneous subpopulations with a direct correspondence to physical single-molecule events, thus offering distinct methodological advantages in resolving sample complexity.

**Fig 4.**
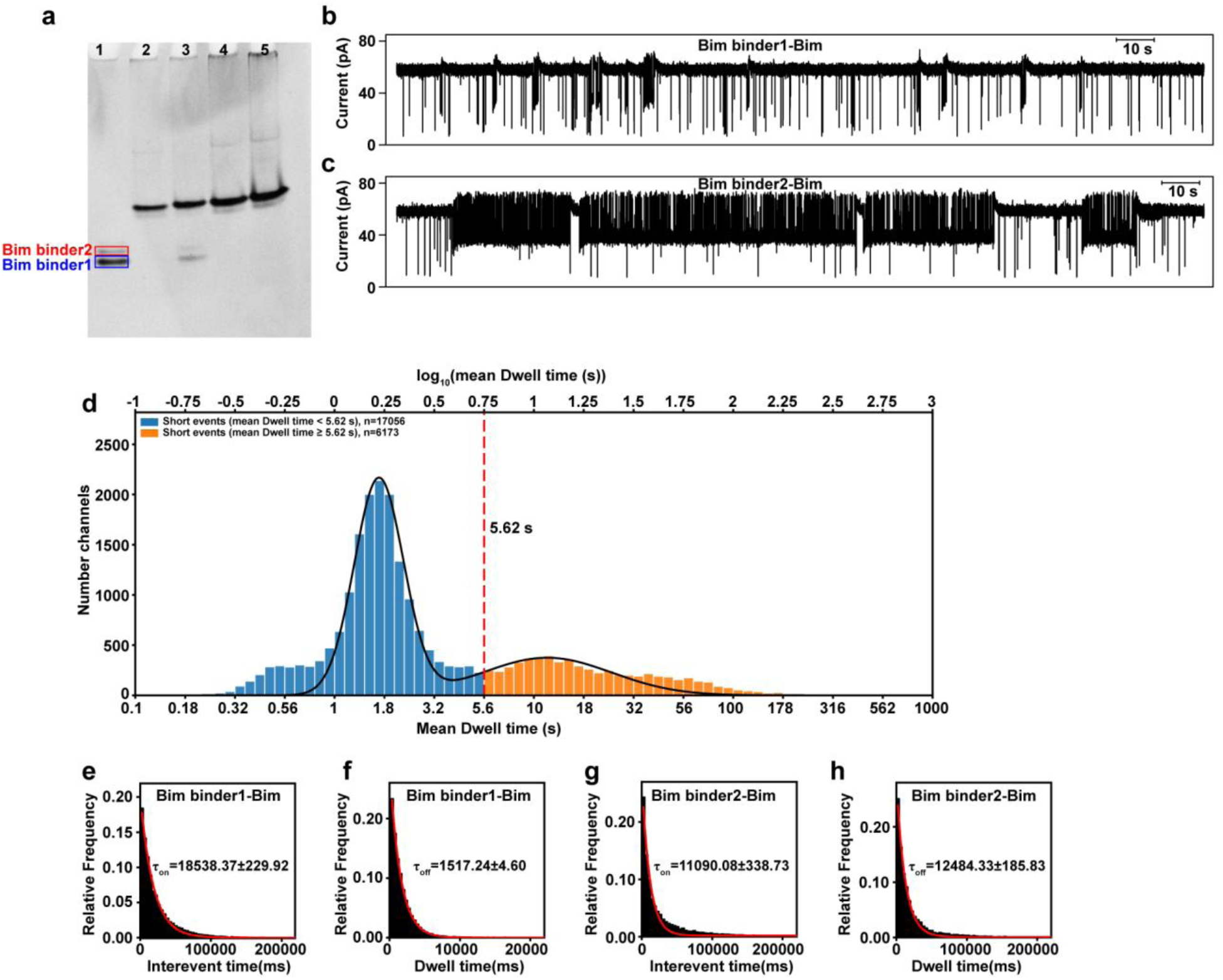
Resolving heterogeneous Bim binder-Bim interactions using DMK. **a,** Native-polyacrylamide gel electrophoresis (native-PAGE) verifying the binding activity of the heterogeneous Bim binder with Bim. Lane 1: Bim binder alone, showing two distinct protein bands. Lane 2: Bim (fused to the C-terminus of the NetB monomer for visualization due to its small size). Lanes 3-5: Bim binder mixed with Bim at molar ratios of 1:1, 1:2, and 1:3, respectively. **b,** Bimodal Gaussian fit of the mean dwell times derived from all functional channels in DMK assays. A threshold of 5.62 s, determined from the statistical distribution, categorizes the active channels into two distinct populations (short and long dwell time groups). **c, d,** Representative current traces for the interaction of Bim binder1 **(c)** and Bim binder2 **(d)** with Bim. e-h, Statistical distributions of interevent times **(e, g)** and dwell times **(f, h)** for the interactions of Bim binder1 **(e, f)** and Bim binder2 **(g, h)** with Bim. The mean interevent time (*τ_on_*) and mean dwell time (*τ_off_*) were obtained by single-exponential fitting. The final concentration of Bim was 50 nM. All experiments were performed in 200 mM KCl, 50 mM phosphate buffer (pH 6.8) under a voltage of ±100 mV.

### 5. Electric-Field Perturbation and Binding Potential Well Robustness

In the DMK system, the potential drop across captured probe complexes could theoretically perturb their dissociation rate via electrostatic forces. Nevertheless, systematic cross-validation against SPR and independent literature data demonstrates consistency in the derived kinetic constants, indicating that with the optimized DMK setup, electric-field-induced kinetic bias is well confined within acceptable thresholds, ensuring the physiological validity of *K_off_* values. Furthermore, for a given molecular pair with fixed conformation and orientation, the applied electric field can be treated as a controllable mechanical parameter. Variable-voltage experiments combined with the Bell-Evans model (Equation 3) allow us to assess the influence of varying electric field strengths on dissociation kinetics and extrapolate the parameters to the zero-field limit^32^, thereby approaching the intrinsic binding characteristics of the molecules free from external forces.

We performed variable-voltage experiments on the CL7-Im7 complex, acquiring kinetic parameters across a voltage range of 60 mV to 140 mV (Fig. 5a-b). According to the Bell-Evans model, ln(*K_off_*) scales linearly with voltage, and the intrinsic *K_off_* value under zero-field conditions can be obtained from the intercept at V=0 via linear extrapolation. For the CL7-Im7 interaction, we observed that *K_off_* exhibits only minor fluctuations across the tested voltage range (Fig. 5c-d), indicating that the complex possesses a deep binding potential well and a small activation displacement (Δχ‡) along the dissociation reaction coordinate. The applied electric field is insufficient to significantly tilt the activation energy barrier within this operating range, suggesting that the interaction is highly resistant to mechanical perturbation.

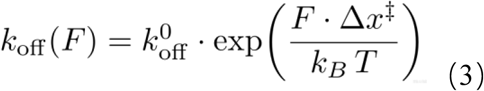

**Fig 5.**
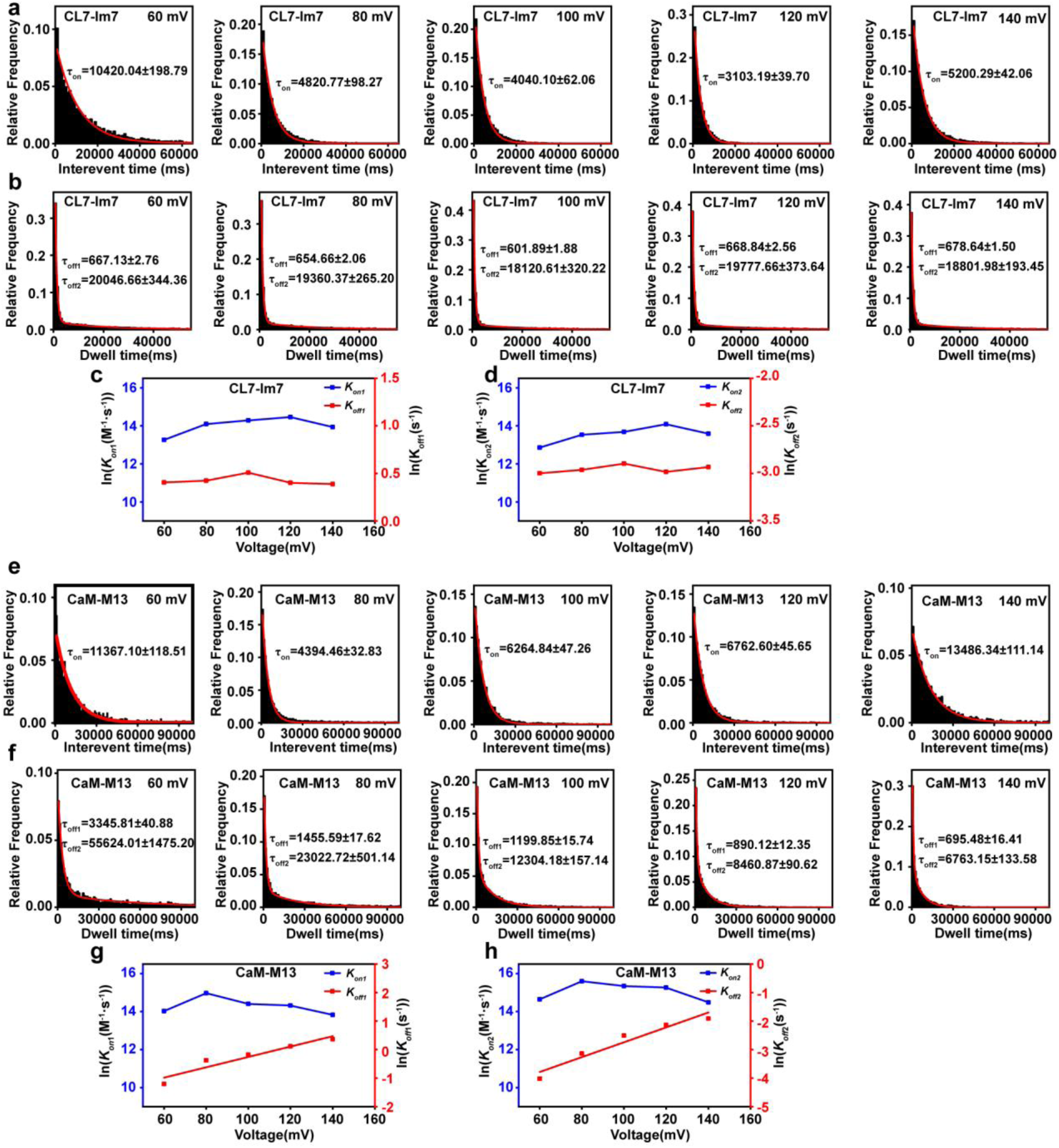
Electric-field perturbation, binding potential well characteristics, and mechanical resistance. **a-d,**Variable-voltage analysis of the interaction between CL7 and 100 nM Im7 across a voltage range of 60 to 140 mV. **a,** Statistical distributions of interevent times at various voltages. **b,** Dwell time distributions, with dwell times fitted by a dual-exponential function to yield *τ_off1_* and *τ_off2_*. **c, d,** Bell-Evans linear regression plots illustrating the voltage sensitivities (slopes) of ln(*K_on_*) and ln(*K_off_*) for subpopulation 1 **(c)** and subpopulation 2 **(d)**, reflecting the activation displacement (Delta_chi_ddagger) and resistance to electric-field perturbation. **e-h,** Variable-voltage analysis of the interaction between CaM and 25 nM M13 (60-140 mV). **e,** Distributions of interevent times. **f,** Distributions of dwell times. **g, h,** Bell-Evans linear regression plots for subpopulation 1 **(g)** and subpopulation 2 **(h)**, where notably steeper slopes correspond to a larger activation displacement (Delta_chi_ddagger) and a shallower binding potential well. All experiments were performed in 200 mM KCl and 50 mM phosphate buffer (pH 6.8), with 50 μM/L CaCl2 supplemented for the CaM-M13 assays.

Following this approach, we also performed voltage-gradient assays on the Calmodulin (CaM)– M13 complex^33^ (Fig. 5e-f). As shown, ln(*K_off_*) of the CaM-M13 pair scales linearly with voltage (Fig. 5g-h), exhibiting a notably steeper slope that corresponds to a larger activation displacement (Δχ‡) in the Bell-Evans model. This indicates that, compared with the CL7-Im7 complex, the CaM-M13 pair features a shallower binding potential well and is more prone to external-force perturbation. In this case, extrapolating the obtained *K_off_* to V=0 yields a value closer to the intrinsic rate (Supplementary Table 8).

It is well established that complex mechanical microenvironments in vivo preclude an unperturbed state^34–36^, highlighting the physiological significance of molecular interaction resilience against environmental perturbation. Comparing the CaM-M13 and CL7-Im7 pairs, although the former exhibits a longer "intrinsic" binding residence time, its shallower potential well means that its actual residence time in complex physiological environments should be cautiously revisited. The DMK various-voltage approach presented here not only mitigates electric-field interference to recover intrinsic binding kinetics, but also provides a convenient method for assessing force-sensitive energy barrier properties, thereby offering multi-dimensional reference value for pharmacology and the pharmaceutical industry.

### 6. Expanded Kinetic Windows and the Resolution of Dissociation Heterogeneity

Ultra-long dissociation analysis has been a challenge for SPR due to baseline drift and insufficient dissociation. Detection of ultra-fast binding events is also limited by mass transport and early-phase kinetic unreliability. The DMK platform achieves a significant extension of the kinetic detection time window. By tracking single-molecule events, the platform eliminates dependence on continuous perfusion and thus is free from baseline drift. Furthermore, mathematically, single-molecule event counting combined with censored fitting resolves the challenge of low dissociation fractions, enabling more accurate kinetic determination. With this foundation, we successfully characterized the binding kinetics between green fluorescent protein (GFP) and its high-affinity nanobody^37^, as illustrated in Fig. 6a-b. Notably, a dissociation residence time exceeding 3 hours posed no limitation to the DMK platform. Given the robustness of our operating principles and experimental workflow, we anticipate that the DMK system can handle molecular pairs with even longer residence times. Regarding rapid transient interactions, we have previously demonstrated the precise kinetic resolution of the Sso7d-DNA pair. Additionally, we successfully resolved the binding kinetics between DNA-polymerase complexes and their complementary nucleotides in the presence of non-catalytic metal ions (Supplementary Fig. 8), demonstrating DMK’s broad applicability and robustness in addressing transient binding scenarios.

**Fig 6.**
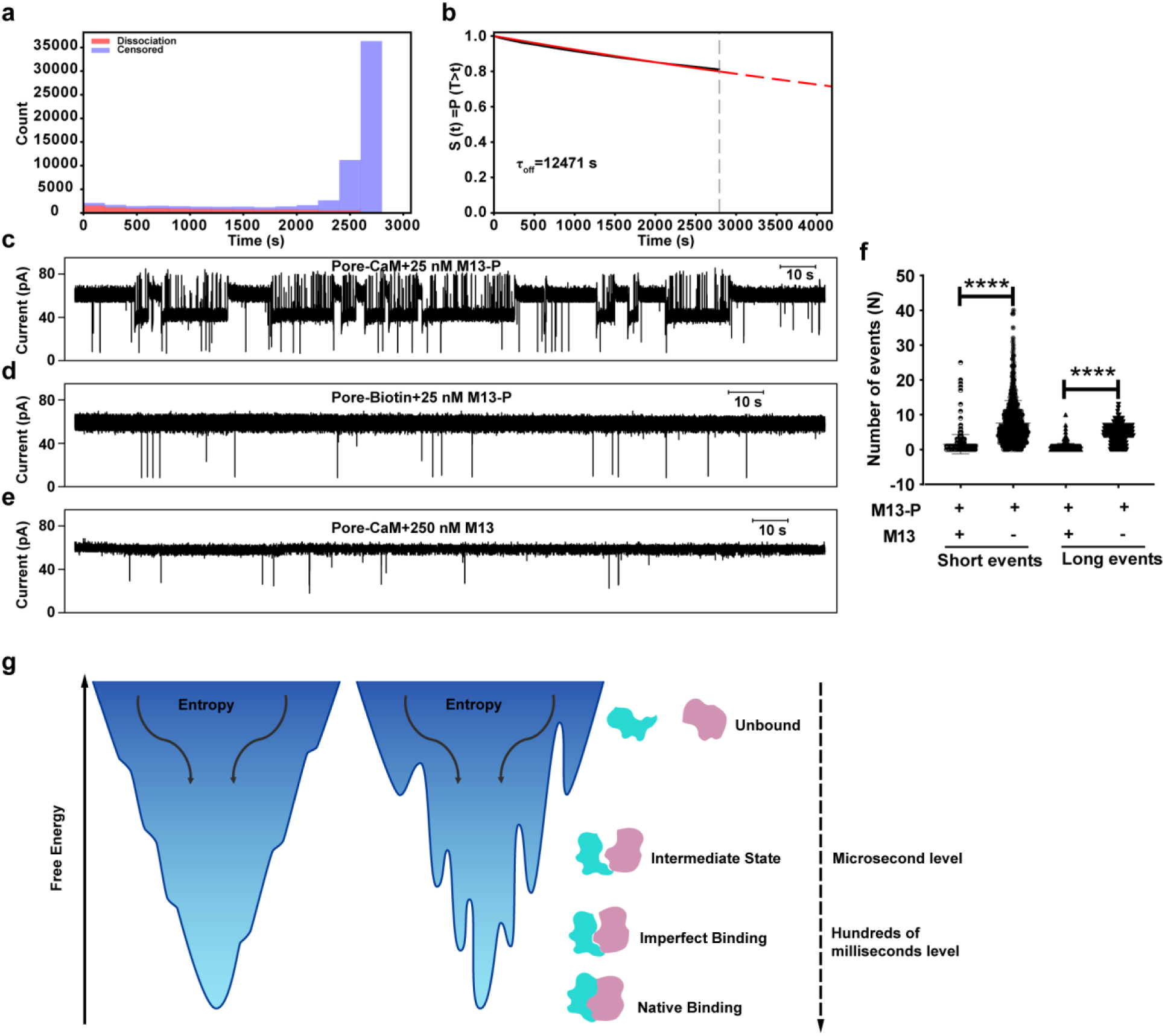
Extension of detection time windows and the imperfect binding model. **a,** Event and censored distribution of dwell times for the interaction between GFP nanobody and 0.195 nM GFP. Red bars represent fully recorded dwell times (11542 events), and blue bars represent censored dwell times (55228 events) for molecules un-dissociated within the recording window. **b,** Maximum likelihood estimation combining complete and censored dwell times to yield the mean dwell time (*τ_off_*). The solid black line (Kaplan-Meier curve) is data-driven without model assumptions, reflecting the true fraction of undissociated molecules within the observation window (plateauing around 0.8). The solid red line shows the single-exponential fit within the observation range, and the dashed red line indicates the extrapolated trend beyond data max. **c–e,** Representative current traces for specificity and competition assays: (**c**) Functional Pore-CaM interacting with 25 nM M13-probe (M13-P, which generates detectable binding signals); (**d**) Negative control using Pore-Biotin interacting with 25 nM M13-P; and (**e**) Competitive control using Pore-CaM interacting with 250 nM un-probed M13 (where M13 binds to CaM but lacks the probe, yielding no nanopore signal and confirming the specificity of the probed interaction). **f,** Quantification of event numbers for short and long dwell times in the Pore-CaM + 25 nM M13-P system upon competition with a 10-fold excess of unlabeled M13 (derived from 500 high-quality, functional channels each). The threshold for short and long dwell times was set to 5 times the fast *τ_off1_* value obtained from dual-exponential fitting of CaM-M13. Both short and long dwell events significantly decreased following the addition of excess M13. Data are shown as mean ± SD. Statistical significance was assessed by unpaired Student’s t-test; **** *p* < 0.0001. All experiments were performed in 200 mM KCl and 50 mM phosphate buffer (pH 6.8). **g,** Energy landscape funnels illustrating different binding modes. Left: smooth energy funnel characterized by a downhill path free of significant energy barriers leading directly to a global free-energy minimum (native binding). Right: rough energy funnel containing multiple metastable states, where molecules starting from the unbound state transition through microsecond-level intermediates before falling into imperfect (non-native, metastable) binding states—corresponding to a residence time of hundreds of milliseconds or above—or reaching the global free-energy minimum (native binding).

Notably, fitting the CaM-M13 dissociation kinetics with a mono-exponential model produced systematic residuals and tailing effects, indicating underlying dynamic heterogeneity. In contrast, a biexponential fitting model resolved two distinct dissociation subpopulations: *τ_off_*_1_, fast (fast phase, ∼1120 ms) and *τ_off_*_2_, slow (slow phase, ∼12304 ms), increasing the coefficient of determination (R^2^) from ∼0.932 to above 0.998 (Supplementary Fig. 9a-b). More importantly, the mathematical biexponential distribution aligns with the physical events captured at the single-molecule level. The obtained slow-phase dissociation parameter (extrapolated to V=0) agreed closely with conventional SPR results. However, the fast phase kinetics derived from rapid dissociation events lacked counterparts in traditional SPR responses. This raised a critical question: do these detected abundant rapid events represent genuine transient specific binding, or are they system artifacts arising from instrumental background or nonspecific interactions?

To address this question, rigorous control experiments were performed. In the negative control assays where the Partner A was not capable of binding to Partner B, or there was no probe on Partner B while all other procedures and components remained identical, both long-and short-residence-time signals were absent (Fig. 6c-e). Furthermore, competitive inhibition assays via the addition of excess free untagged M13 showed synchronous suppression of both signal populations (Fig. 6f). These controls support the interpretation that the fast-phase events originate from specific intermolecular binding.

This dissociation heterogeneity is not unique to the CaM-M13 system. Similar biexponential fast and slow phases were observed across diverse molecular pairs, including CL7-Im7 and CL7_M-Im7 demonstrating broad universality and arguing against preparation-induced heterogeneity of specific ligands. Furthermore, the time constant τ of the fast phase exhibits a strong dependence on the specific molecular pair, ranging from hundreds of milliseconds for CaM-M13 to over 20 seconds for the CL7_M-Im7 high-affinity mutant (Supplementary Fig. 9c-d). Together with the observation that the mSA-biotin pair—characterized by a dissociation time of up to 120 seconds—conforms to a standard mono-exponential distribution without observable heterogeneity, multi-dimensional evidence argues against any instrumental or algorithmic artifacts while supporting that the fast-phase events represent authentic, molecular-pair-dependent kinetic trajectories.

Conventional techniques such as SPR struggle to capture transient binding events with sub-second or shorter *τ_off_* values; even when *τ_off_* values reach 20 seconds and fall within the operational window of SPR, lower-affinity events are often masked by high-affinity, slow-dissociating populations. This limitation has fostered an oversimplified unbound-versus-bound binary view of binding kinetics. Rather, the pathway of molecular interactions along the energy landscape funnel is rarely smooth, particularly for macromolecules involving conformational changes and induced fit, as well as the formation or rearrangement of salt bridges, hydrogen bonds, hydrophobic interactions, and van der Waals forces^38^. These complex processes generate local minima along the funnel, representing various binding intermediates. Previous molecular simulations have identified microsecond-to-millisecond transient intermediates^39,40^. Although current computational limitations restrict simulations to shorter timescales, preventing the capture of intermediates on much longer timescales, the underlying physical principles are consistent with our observations. We thus propose that local minima with energy fluctuations also exist near the bottom of the energy landscape funnel, corresponding to imperfect binding states (Fig. 6g). These states may either surmount activation energy barriers to reach the optimal bound configuration or dissociate rapidly, directly corresponding to the fast-binding events resolved by DMK.

## Discussion

Although applying nanopores to interaction kinetics has long been explored, traditional approaches lack broad universality due to analyte-dependent charge and structural constraints. While probe-labeling expands the target scope, confining probes near the channel restricts detection to binding-induced conformational changes^18,41^. This dependency requires extensive trial-and-error, trapping methods in a "one-pore, one-target, one-probe" customization bottleneck. In contrast, the present platform labels probes on free solution molecules, exploiting binding-induced local concentration enhancements to bypass structural constraints for universal detection. Additionally, high-frequency AC filtering eliminates the background interference and pore-clogging associated with DC schemes. Moreover, pre-engineered click chemistry handles simplify target immobilization to a standard procedure akin to streptavidin-mediated surface capture in SPR. These breakthroughs establish a versatile, universal platform capable of successfully resolving interaction kinetics across diverse molecular pairs—encompassing protein, peptide, DNA, and small-molecule systems—with high concordance to established SPR and literature benchmarks. Moving beyond macroscopic signal analysis, the DMK system harnesses single-molecule event-based resolution and achieves an expanded kinetic window, overcoming conventional limits in resolving transient binding, heterogeneity, and complex kinetics. Furthermore, integrated thermal and electrical modulation enables multidimensional interaction profiling. Specifically, variable-temperature assays yield ITC-comparable thermodynamics while enabling direct kinetic characterization at physiological temperature. Concurrently, voltage-gradient assays reveal distinct responses to electric-field stress among pairs with comparable apparent kinetics, providing additional insights for assessing interaction stability.

DMK analyses revealed prevalent heterogeneity in dissociation kinetics across several molecular pairs, comprising both fast and slow components. A retrospective examination of raw SPR data reveals residual long-tail effects under single-exponential fitting, driven by rapid signal decline in early dissociation stages. While switching to a double-exponential fit improves fitting quality (Supplementary Fig. 10), SPR is constrained by coupling interference from buffer-exchange perturbations and non-specific adsorption. Consequently, traditional analyses dismissed this initial phase as an experimental artifact, retaining only the dominant slow-dissociation component corresponding to long residence times. In contrast, the fast-dissociation components in the DMK dataset correspond directly to single-molecule events, allowing for a direct verification of this heterogeneous dissociation by negative controls and competition assays. Furthermore, the observation of dynamic heterogeneity across multiple molecular pairs exhibiting a broad range of pair-specific τ_off_ values, together with the homogeneous control of the mSA-biotin interaction, provides additional evidence against instrumentation or algorithmic artifacts. Interpreting the fast dissociation events captured by DMK as ’imperfect binding’ aligns with short-timescale molecular dynamics simulations and local minima on the energy landscape, providing a complementary view of molecular interaction complexity. Traditionally, pharmacology and drug discovery have emphasized long residence times. However, overlooking short-residence-time events (accounting for 28% to 65% of DMK-detected interactions) potentially creates a blind spot in biological understanding. Despite their sub-second or hundred-millisecond duration, these transient events can drive profound biochemical effects in core processes like signal transduction^42^, prompting a re-evaluation of traditional paradigms. Nevertheless, as a technology development team primarily focused on platform innovation, our scope is limited in exploring intricate biological or pharmacological mechanisms. We look forward to future cross-validation by researchers in the field using alternative platforms (e.g., smFRET), broader generality analyses across diverse molecular pairs, and in-depth explorations of the functional impacts of such heterogeneous dissociation on biochemical and pharmacological outcomes, thereby advancing our understanding of molecular interaction theories and facilitating biochemical applications.

Like any analytical platform, DMK has intrinsic limitations and boundaries. Because measurement inevitably perturbs the analyte, DMK data reflect dynamic approximations under specific conditions (e.g., electric fields and probe labeling) rather than absolute ground truth. Electric fields are intrinsic to nanopore sensing and the DMK workflow. The high concordance of our data with SPR and literature benchmarks indicates that our optimized designs effectively constrain electric field perturbations within an acceptable range. For interactions sensitive to electric fields, voltage-gradient assays provide a convenient way to evaluate such perturbations and even assess binding resilience in complex physiological environments. However, given the intricate multi-factor coupling within the system—such as molecular conformation, orientation, and probe-nanopore electrical response—these voltage-gradient results currently serve primarily as a qualitative reference rather than a definitive basis for absolute quantitative analysis. Future investigations incorporating tailored coupling geometries, probe designs, and channel features will yield deeper mechanical insights into intermolecular interactions, thereby advancing our understanding of molecular interaction theories and facilitating biochemical applications. Regarding probe labeling, it is a standard biochemical auxiliary widely applied across diverse research contexts. For the vast majority of proteins, peptides, or nucleic acids, terminal-labeling-induced perturbations are generally negligible. For small molecules, however, labeling-induced deviations are more prone to occur. Nevertheless, such deviations can be evaluated and corrected through competition assays using unmodified natural counterparts on the platform. Taken together, despite the practical boundaries, DMK provides a reliable and robust framework for kinetic characterization.

Compared with SPR, DMK broadens application scope but introduces an extra probe-labeling step in addition to the nanopore-coupling procedure (which is analogous to SPR chip immobilization). For the labeling of diverse molecular types—including proteins, peptides, nucleic acids, and various functional groups of small molecules—modern biochemistry provides established modular modification schemes. This enables the straightforward preparation of probe-labeled molecules, granting the platform the operational convenience required for large-scale deployment. Moreover, drawing upon a mature hardware architecture already scaled for commercial application, the high-throughput nanopore array provides exceptional system robustness. Even with yield fluctuations in active pores caused by various factors, the system routinely delivers over 10,000 qualified channels, enabling the parallel acquisition of statistically robust, massive single-molecule event data. More importantly, this platform shares an identical hardware architecture and chip system with single-molecule sequencing systems^43–45^, allowing researchers to seamlessly switch between sequencing and kinetic assays simply by changing the reagent kit and analysis software. This multifunctional integration offers high ease of deployment and maintenance. Combined with the inherent electrical cost advantages of nanopore technology, the DMK system provides a versatile, accessible, and cost-effective framework, holding substantial promise for breakthroughs in modern life sciences and applied research.

## Methods

### Protein Expression

The target gene encoding a 6xHis-tagged protein was cloned into the pET-26b plasmid, and the sequence-verified plasmid was transformed into E. coli ER2566 competent cells. A single colony was inoculated into LB medium supplemented with 50 ug/mL kanamycin and grown at 37 °C with shaking at 250 rpm until the OD600 reached 0.8. Protein expression was then induced by adding 0.2 mM isopropyl-beta-D-thiogalactoside (IPTG), followed by incubation at 16 °C and 250 rpm for 20 h. Cells were harvested by centrifugation at 2851 *g* for 15 min at 4 °C.

### Purification of Nanopore Monomers and Im7/GFP

In this study, the nanopores we used were the NetB proteins from Clostridium perfringens toxin^46^. Cells were resuspended in lysis buffer (pH 7.0, 50 mM sodium phosphate, 200 mM NaCl, 10% glycerol) and lysed by sonication. The lysate was centrifuged at 24149 *g* for 15 min, and the supernatant was collected and loaded onto a Ni-Bestarose FF resin column for affinity purification. The column was washed with wash buffer (pH 7.0, 50 mM sodium phosphate, 200 mM NaCl, 10% glycerol, 30 mM imidazole) to remove impurities, and the target protein was eluted using elution buffer (pH 7.0, 50 mM sodium phosphate, 200 mM NaCl, 10% glycerol, 300 mM imidazole). The eluted protein was further purified by size-exclusion chromatography using a Superdex 200 Increase 10/300 GL column equilibrated with storage buffer (pH 7.0, 50 mM sodium phosphate, 200 mM NaCl, 10% glycerol) to yield the purified nanopore monomers or Im7/GFP proteins.

### Purification of mSA

As the mSA protein was expressed as inclusion bodies, the harvested cells were resuspended and lysed in a buffer containing 6 M guanidine hydrochloride by sonication. Following high-speed centrifugation, the supernatant was loaded onto a Ni-Bestarose FF resin column. Impurities were removed using a wash buffer containing 6 M guanidine hydrochloride, and the denatured mSA protein was eluted using an elution buffer containing 6 M guanidine hydrochloride. To refold the protein, the denatured mSA was added dropwise into an ice-cold refolding solution (50 mM sodium phosphate, 150 mM NaCl, 0.3 mg/mL biotin, 0.2 mg/mL oxidized glutathione, and 1 mg/mL reduced glutathione, pH 7.0) with rapid stirring. The refolded protein was subjected to a second round of affinity purification using Ni-Bestarose FF resin, washed with wash buffer (pH 7.0, 50 mM sodium phosphate, 200 mM NaCl, 10% glycerol, 30 mM imidazole) to remove impurities, and eluted with elution buffer (pH 7.0, 50 mM sodium phosphate, 200 mM NaCl, 10% glycerol, 300 mM imidazole). Finally, the protein was further purified by size-exclusion chromatography using a Superdex 200 Increase 10/300 GL column equilibrated with storage buffer (pH 7.0, 50 mM sodium phosphate, 200 mM NaCl, 10% glycerol) to yield the purified mSA protein.

### Purification of DNA polymerase^47^ and Sso7d

Cells were resuspended in lysis buffer (pH 7.0, 50 mM sodium phosphate, 1 M NaCl, 10% glycerol, 2 mM TCEP) and lysed by sonication. Following high-speed centrifugation, the supernatant was loaded onto a Ni-Bestarose FF resin column for affinity purification. The column was washed with wash buffer (pH 7.0, 50 mM sodium phosphate, 1 M NaCl, 10% glycerol, 2 mM TCEP, 30 mM imidazole) to remove impurities, and the target protein was eluted using elution buffer (pH 7.0, 50 mM sodium phosphate, 200 mM NaCl, 10% glycerol, 2 mM TCEP, 300 mM imidazole). The eluted protein was further purified using a HiTrap Heparin column with isocratic salt elution using 800 mM NaCl in 20 mM sodium phosphate (pH 7.0), followed by size-exclusion chromatography on a Superdex 200 Increase 10/300 GL column equilibrated with storage buffer (pH 7.0, 50 mM sodium phosphate, 200 mM NaCl, 10% glycerol) to yield the purified DNA polymerase or Sso7d protein.

### Fusion Expression of the Nanopore Monomer and Partner A Protein

The full-length coding sequences of CL7/SpyCatcher/Bim Binder were fused to the C-terminus of the nanopore monomer protein via a flexible Gly-Ser (GS) linker, and the resulting tandem fusion construct was co-expressed. The purification procedure followed the same protocol as that described for the nanopore monomer.

### Preparation of Heteroheptameric Nanopore Proteins (6:1 Stoichiometry)

The preparation of heteroheptameric nanopores was performed following previously established methods [Design and characterization of a nanopore-coupled polymerase for single-molecule DNA sequencing by synthesis on an electrode array]. Briefly, the nanopore monomer and the fusion monomer protein (e.g., nanopore monomer-CL7/SpyCatcher/Bim Binder) were mixed at a molar ratio of 5:1. The lipid 1,2-diphytanoyl-sn-glycero-3-phosphocholine (DPhPC) was added to a final concentration of 1 mg/mL, followed by incubation at 37 °C for 2 h. After the addition of 2.5% beta-OG and 0.2% Tween-20, the mixture was centrifuged at 2851 *g* for 3 min. The supernatant was then loaded onto a Resource Q ion-exchange column and purified using a linear salt gradient from 20 mM to 2 M NaCl in 20 mM sodium phosphate and 0.2% Tween-20 (pH 7.0). The desired 6:1 assembly eluted after the homoheptameric (7:0) nanopore protein because the 6:1 assembly contained nineteen engineered Asp-Glu tags (19x-Asp-Glu-tag), yielding the Partner A-bearing 6:1 heteroheptameric nanopore protein.

### Chemical Coupling of Partner A to Nanopore

In this study, mSA, GFP nanobody, CaM, or DNA polymerase was chemically coupled to nanopores. Briefly, an excess of I-PEG3-N3 (Bidepharm, CAS 936917-36-1) was added to cysteine-mutated nanopore monomers. Following incubation at room temperature for 2 h, the mixture was purified using a Resource Q ion-exchange column with a linear salt gradient from 20 mM to 2 M NaCl in 20 mM sodium phosphate (pH 7.0) to yield N3-modified nanopore monomers. Unmodified nanopore monomers and N3-monomers were then mixed at a molar ratio of 5:1 with 1 mg/mL phospholipids, followed by incubation at 37 °C for 2 h. After the addition of 2.5% beta-OG and 0.2% Tween-20, the mixture was centrifuged at 2851 *g* for 3 min. The supernatant was treated with TEV protease to remove the His-tag from the nanopore proteins at room temperature for 2 h, and subsequently purified using a Resource Q ion-exchange column with a linear salt gradient from 20 mM to 2 M NaCl in 20 mM sodium phosphate and 0.2% Tween-20 (pH 7.0) to yield His-tag-free, N3-modified 6:1 nanopores. Pre-engineered nanopore reagents containing the N3 handle can be purchased directly from Geneus (Geneus, 036.001.001).

Partner A was mixed with NHS-ester-PEG3-DBCO (Confluore, CAS 2163772-16-3) at a 1:1 molar ratio and reacted at room temperature for 2 h in reaction buffer (pH 6.8, 50 mM sodium phosphate, 200 mM NaCl). Unreacted NHS-ester-PEG3-DBCO was removed by washing during purification on a Ni-Bestarose FF resin column. The eluted DBCO-modified Partner A was subsequently reacted with N3-functionalized 6:1 heteroheptameric nanopores at room temperature for 2 h via copper-free click chemistry. The reaction mixture was loaded onto a Ni-Bestarose FF resin column, washed with wash buffer (pH 7.0, 50 mM sodium phosphate, 200 mM NaCl, 10% glycerol, 0.2% Tween-20, 30 mM imidazole) to remove unreacted nanopores, and eluted with elution buffer (pH 7.0, 50 mM sodium phosphate, 200 mM NaCl, 10% glycerol, 0.2% Tween-20, 300 mM imidazole). Because the 6:1 assembly retained six His-tags, the target Partner A-conjugated 6:1 heteroheptameric nanopore was selectively eluted and recovered.

For chemical coupling of biotin to nanopores, an excess of Maleimide-PEG6-biotin (TCI, CAS 1808990-66-0) was added to cysteine-mutated nanopore monomers. Following incubation at room temperature for 2 h, mass spectrometry confirmed 100% modification stoichiometry. The biotin-modified 6:1 heteroheptameric nanopores were subsequently prepared following the established heteroheptameric nanopore assembly protocol.

For chemical coupling of DNA to nanopores, the PEG6-DNA sequence (5’-GTGCAGTTCACGGATGCTGTT-3’) was synthesized via solid-phase synthesis using a single-stranded DNA synthesizer (12P, Beijing Tsingke Biotech). An acetonitrile solution of 4-maleimidobutyric acid N-hydroxysuccinimide ester was added to the upper aqueous phase of the reconstituted DNA solution (pH 7.0). After reaction at room temperature for 1 h, Maleimide-PEG6-DNA was purified by HPLC. An excess of Maleimide-PEG6-DNA was then added to cysteine-mutated nanopore monomers and allowed to react at room temperature for 2 h. The reaction mixture was purified using a Resource Q ion-exchange column with a linear salt gradient from 20 mM to 2 M NaCl in 20 mM sodium phosphate (pH 7.0) to yield DNA-modified nanopore monomers. Finally, the DNA-modified 6:1 heteroheptameric nanopores were prepared following the standard heteroheptameric assembly procedure.

### Probe Reagents for Coupling

N3-functionalized probe reagents for coupling can be purchased directly from Geneus (Geneus, 036.003.001).

### Chemical Coupling of Probe to Partner B

In this study, the probe was coupled to the N-terminus of Im7, mSA, or Sso7d proteins. Purified proteins were mixed with NHS-PEG3-DBCO at a 1:1 molar ratio and reacted in a reaction buffer (pH 6.8, 50 mM sodium phosphate, 200 mM NaCl) at room temperature for 2 h. Unreacted NHS-ester-PEG3-DBCO was removed by washing during purification on a Ni-Bestarose FF resin column. The eluted DBCO-modified proteins were subsequently reacted with an excess of the N3-functionalized probe via copper-free click chemistry at room temperature for 2 h. The reaction mixture was loaded onto a Ni-Bestarose FF resin column to remove excess unreacted probes, and further purified using a Resource S ion-exchange column with a linear salt gradient from 20 mM to 2 M NaCl in 20 mM sodium phosphate (pH 7.0) to yield probe-conjugated Partner B. For the chemical coupling of small-molecule biotin to the probe, DBCO-biotin (Confluore, CAS 1418217-95-4) was reacted with the N3-functionalized probe via click chemistry. The reaction product was purified by high-performance liquid chromatography (HPLC) on a C18 reverse-phase column using a linear gradient of 5% to 40% acetonitrile in 1% formic acid aqueous solution over 30 min to yield the probe-functionalized small molecule.

### Fusion Expression of Probe-Partner B

In this study, SpyTag, M13, or Bim was genetically fused with the probe to replace chemical peptide synthesis and modification. The fusion construct was designed in a tandem configuration as 6xHis-tag-green fluorescent protein (GFP)-TEV cleavage site-SpyTag/M13/Bim-probe. Following expression, the proteins were purified using a Ni-Bestarose FF resin column. Imidazole was removed using a G25 desalting column, and the sample was then incubated with TEV protease for 2 h to cleave the 6xHis-GFP tag. The mixture was passed through a Ni-Bestarose FF resin column again to collect the flow-through, which was further purified by size-exclusion chromatography on a Superdex 200 Increase 10/300 GL column (buffer: pH 7.0, 50 mM sodium phosphate, 200 mM NaCl, 10% glycerol) to yield probe-fused SpyTag, M13, or Bim.

### Protein Purity and Activity Validation

The purity of all purified proteins was assessed by sodium dodecyl-sulfate polyacrylamide gel electrophoresis (SDS-PAGE). Protein activity was verified through intermolecular binding assays using native polyacrylamide gel electrophoresis (native-PAGE).

### DNA polymerase Activity Assay

The 5’-Cy5-modified hairpin nucleic acid and the 3’-BHQ2-modified complementary nucleic acid were mixed at a molar ratio of 1:1.2, heat-treated at 95°C for 5 min, and naturally cooled to anneal, forming a double-stranded hairpin nucleic acid complex. An aliquot of 25 nM annealed nucleic acid was then incubated on ice for 10 min with the enzyme at an enzyme-to-nucleic acid molar ratio of 4:1. Subsequently, potassium acetate (KAc at a final concentration of 650 mM), Tris (pH 7.5, 20 mM), 5% ethylene glycol, and various metal ions were added, with water added to bring the total volume to 100 μL. The reaction was performed in a 96-well microplate (Black 96-well cell culture plate, Flat Bottom, WHB). A final concentration of 5 μM dNTP was injected via the automated injector of a microplate reader (Synergy H1, Bio-Tek), and detection was carried out at an excitation wavelength of 540 nm and an emission wavelength of 570 nm.

### Preparation of Nanopore-Polymerase-DNA ternary complex

The hairpin DNA was heat-treated at 95°C for 5 min and then allowed to cool naturally to anneal into double-stranded hairpin nucleic acids. An aliquot of 1 μM annealed nucleic acid was then incubated with the DNA polymerase-nanopore complex at a molar ratio of 20:1 (nucleic acid to complex) at 25°C for 1 h to assemble the DNA-polymerase-nanopore complex.

### High-Throughput Nanopore Sensing

Molecular interaction assays were performed using the G-seq500 instrument and G-MK02 chips (Geneus Technologies). Lipid membrane formation and the insertion of nanopore-partner A molecules were carried out automatically based on the platform’s built-in pump-valve system and protocol workflow, using a nanopore-partner A concentration of 100 pM in a 100 μL volume. The software automatically filters out invalid channels or channels with multiple nanopore insertions based on open-channel current amplitudes, recording only normal single-channel pores. Due to variations in conjugation efficiency and purity, the number of active channels per experiment may fluctuate significantly, but typically over 10,000 active channels are achieved. The acquired electrical signals are digitized and noise-filtered before being automatically saved for downstream analysis. Unless otherwise specified, all measurements were performed at 25°C.

### Kinetic analysis based on macroscopic association-dissociation curves

The impedance range of single open pores was determined based on channel impedance statistics. For kinetic analysis based on macroscopic impedance signals, all channels identified by the software as exhibiting normal open-pore currents (impedances) were included in the statistics, yielding active channel numbers typically ranging from 10,000 to over 100,000 for different molecular pairs. For binding assays, Partner B bearing the probe was perfused into the cis chamber. Unless otherwise specified, a brief alternating voltage of ±100 mV was applied to each channel, and the chip’s active channel impedance was calculated every 4 s (consisting of 4 electrical cycles, with 40 ms for positive and 40 ms for negative polarization per cycle). The numbers of open pores (unbound) and blocked pores (bound) were determined and recorded in real time. For dissociation assays, the buffer was continuously perfused through the chip. Custom event detection was implemented using MATLAB (2018b; MathWorks). Reaction rates (blocking ratios) were calculated based on the numbers of open and blocked channels: blocking ratio = blocked channels / (blocked channels + open channels). The resulting blocking ratio-versus-time curves yielded the association/dissociation-versus-time curves. Statistical data were analyzed using GraphPad Prism 8.0, and curves were fitted using nonlinear regression programs (Association and dissociation) to calculate *K_on_*, *K_off_*, and *K_d_*, or fitted via association kinetics to calculate *K_on_* For molecular pairs requiring steady-state fitting, analysis was performed using the Hill equation to determine *K_d_*.

### DMK Kinetic Analysis Based on Single-Molecule Discrete Events

Unless otherwise specified, single-molecule nanopore recordings were performed using an alternating current with positive and negative polarizations applied for 40 ms each. The collected current signals from each channel were subjected to density-based clustering analysis on the time-series data within individual channels. For each sampling point, the current difference between it and subsequent sampling points was evaluated to determine whether adjacent sampling points shared similar current levels, thereby grouping consecutive sampling points with similar current features into identical data clusters. Based on the current distribution features of each cluster, the state boundary between the blocked state and the open state was established, enabling state annotation along the continuous time series. This allowed the determination of the dwell time of blockade events and the interevent time between consecutive blockade events.

To ensure that recorded events correspond to genuine binding signals rather than stochastic, transient captures of free molecules across voltage cycles, a valid event was required to be detected in 3 consecutive cycles. Specifically, for an alternating waveform of ±40 ms per phase (80 ms per full cycle), an approximate dwell-time cutoff of 160 ms was applied (corresponding to events spanning at least two full cycles). Events with dwell times below this threshold were excluded from final fitting due to the system’s time-resolution limits. For interactions with a *τ_off_* significantly longer than the time resolution (exceeding 10-fold), the proportion of missed events below 160 ms was kept within 10%, rendering its impact on *K_on_* negligible. For Koff fitting, a truncated fitting approach was employed to circumvent the bias introduced by missed short events. For rapid dissociation events where *τ_off_* approached the time-resolution limit, a first-order correction of the on-rate based on maximum likelihood was applied (data not shown), thereby mitigating underestimation of the On-rate caused by system temporal limitations. Conversely, truncated fitting minimized the influence of time resolution on the off-rate. For ultra-slow dissociating molecular pairs (e.g., GFP nanobody-GFP), given that a large fraction of molecules remained in the bound state without dissociating within the measurement window, maximum likelihood estimation was utilized during analysis to prevent errors arising from incomplete signal recording.

Similarly, for generalized kinetic profiling across diverse or unknown molecular pairs—where the exact extent of stochastic signal dropouts caused by fluctuations in capture rate or molecular conformation is difficult to predict a priori—establishing an interval of 5 consecutive cycles without a probe signal provides a robust safety margin. By tuning the concentration of Partner B to keep the mean interevent time *τ_on_* well above 10-fold this threshold, the proportion of missed true intervals can be effectively confined within 10%. Combined with truncated fitting across all relevant kinetic analyses, this strategy converts unpredictable microscopic uncertainties into a controlled error bound, securing broad safety redundancy for universal parameter extraction.

To resolve the ultrafast association and dissociation kinetics of dNTP/metal ion-polymerase/DNA complexes, the AC waveform was adjusted to a 2-ms cycle (1 ms each for the positive and negative phases), with both the dwell-time and interevent interval cutoffs set to 4 ms. It should be noted that such short time cutoffs are not universal parameters across this platform, but rather represent platform-specific optimizations enabled by the exceptionally high capture efficiency of these specific probes. For other molecular pairs in this study without such dedicated optimization, a voltage half-cycle of 40 ms serves as a more reliable choice, as over 90% of probe capture events occur within 10 ms (data not shown).

In the current trace displays, except for representative schematics that retain complete positive and negative voltage cycles, the remaining comparative traces omit the reverse voltage segments to facilitate intuitive comparison, presenting only concatenated consecutive forward voltage traces. It should be noted that all temporal statistics and kinetic calculations remain strictly based on the full alternating cycle duration and are unaffected by this visualization format.

Unless otherwise specified, the extracted interevent time and dwell time data were fitted using Origin 2021 via exponential fitting to obtain the mean interevent time (*τ_on_*) and mean dwell time (*τ_off_*). Because the GFP nanobody-GFP pair dissociates extremely slowly, some binding events remained unresolved at the end of the recording window, and their dwell times could not be fully captured. We therefore pooled the completely and incompletely recorded dwell-time data and fitted the combined dataset by maximum likelihood estimation using a script written in Python. The association rate constant (*K_on_*) was determined using the formula *K_on_* = 1/ (*τ_on_* * [C]), where [C] represents the concentration of the analyte added to the cis solution, and values for the dissociation rate constant (*K_off_*) were determined as 1 / *τ_off_*. The equilibrium dissociation constant (*K_d_*) was calculated as *K_off_* /*K_on_*. In the thermodynamic calculations, *K_d_* values obtained at different temperatures were used to further calculate the association equilibrium constant (*K_A_* = 1/*K_d_*) for subsequent thermodynamic parameter determinations. In the exponential fitting analysis, the dissociation processes of DNA-Sso7d, Biotin-mSA, and Bim binder-Bim well conformed to a single-exponential distribution, whereas the dissociation process of CaM-M13 exhibited pronounced heterogeneity. As control experiments thoroughly confirmed the authenticity of the fast-phase dissociation events rather than system noise, a biexponential fitting model was adopted for analysis, and the kinetic parameters for State 1 and State 2 were resolved and calculated based on the relative amplitudes of the two phases. Another molecular pair exhibiting a heterogeneous distribution in dissociation was CL7_M-Im7, for which a biexponential fitting model was also employed. Although the CL7-Im7 molecular pair likewise displayed heterogeneous distribution features, a single-exponential fitting model was applied to homogenize the fast and slow phases to a certain extent for approximation, enabling direct comparison with the ensemble macroscopic data from isothermal titration calorimetry (ITC).

Because the identification and extraction of single-molecule events impose higher requirement on signal quality, the algorithm assigns a Q-score to each channel for single-molecule kinetic analysis, similar to Q-score prediction in sequencing, based on parameters such as open-pore current amplitude, baseline stability, overall channel noise level, signal capture rate, and blocking current magnitude. For molecular pairs with short signal durations (dNTP/metal ions-polymerase/DNA), each channel can generate hundreds of events, allowing kinetic results to be obtained through statistical analysis of a single channel. Furthermore, pooling events from hundreds of high-Q-score channels yields higher-throughput statistical results. For molecular pairs with longer signal durations, high-Q-score channels are selected, and channels with obviously inaccurate Q-score assessments (approximately 5%) are filtered out via manual review; statistical analysis of thousands of pooled events is then performed to obtain the kinetic results. For heterogeneous sample (Bim binder) analysis, the mean dwell time of all normal channels was calculated, and a threshold was established via bimodal Gaussian fitting of its statistical distribution plot, thereby enabling classification and independent kinetic analysis of channels corresponding to different subpopulations.

### Surface Plasmon Resonance (SPR) Kinetic Assay and Analysis

Binding assays for molecular pairs A and B were performed using a Biacore 1K+ system (Cytiva). Featuring a single-needle, six-flow-cell configuration and equipped with a Series S Sensor Chip CM5, the instrument was used to covalently immobilize ligand protein A onto the chip surface via standard amine coupling. Sample injection and data acquisition were controlled by the accompanying Biacore Insight software. Except for Bim, which is a probe-bearing sample (owing to the relatively small size of the peptide), all other analytes were unmodified free molecules. Measurements were conducted across multiple concentrations using the single-cycle kinetics (SCK) mode. After subtracting signals from the reference flow cell and blank cycles using the Biacore Insight software, the sensorgrams were globally fitted using a 1:1 Langmuir binding model to extract kinetic parameters. For the heterogeneous Bim binder-Bim molecular pair, a heterogeneous ligand model was further applied for curve fitting.

### ITC Thermodynamic Measurement and Analysis

The ITC measurements were performed as an independent third-party testing service by Ceshigo Research Service Agency using the samples provided by Geneus. The thermodynamic characterization of the interaction between the CL7-Im7 molecular pair was performed using a NANO ITC isothermal titration calorimeter (TA Instruments). All solutions and samples were vacuum-degassed for 10 min prior to use. The CL7 solution (1 μM) was loaded into the sample cell, and the Im7 solution (10 μM) was loaded into the titration syringe. The measurements were conducted at 25 C with 20 successive injections, maintaining an interval of 200 seconds between each injection. The reference cell was filled with buffer. Data were analyzed using TRIOS software (version 6.0, TA Instruments). The peak areas of each injection after baseline correction were integrated to obtain the heat of reaction per injection. The corrected injection heats were fitted to an independent binding site model to yield the equilibrium dissociation constant (*K_d_*), stoichiometry (n), enthalpy (delta H), and entropy (delta S). The change in Gibbs free energy (delta G) was calculated using the equations delta G = -RT ln Ka and delta G = delta H -T delta S.

## Supporting information

SI merterial

## Acknowledgements

We thank Rongsheng Luo, Shu Liu, Lei Li, Xinyuan Huang, Wanhui Jiang, Bo Yan, and Jiguo Hou for their technical support in developing perfusion scripts, modifying software/hardware configurations, and optimizing temperature control systems during data acquisition, and Ke Jiang for providing the Python script for maximum likelihood fitting. We thank Ting Xie for assistance with figure refinement and graphic aesthetics. We also acknowledge Chao Tang and Xu Deng for their contributions to early-stage exploratory work, as well as Ronghuo Qin, Kai Chen, and Jun Huang for helpful discussions and constructive suggestions. We thank Min Chen for contributions to the verification of sample preparation work. This work was supported by Geneus internal R&D funding.

## Author contributions

Y.Z.Z. conceived the project, led the experimental design, supervised data analysis, and wrote the manuscript. K.J.L. contributed to experimental design, performed experiments, analyzed the data, and prepared the manuscript. Y.W. prepared biochemical samples, performed data acquisition, and wrote the Methods section. Y.W.C. developed event extraction algorithms and implemented channel Q-value filtering. T.T.Z. synthesized probes and click chemistry linkers. S.Y.L. led molecular biology work as well as protein expression and purification. L.P.Z. wrote analytical scripts and participated in macro-kinetic curve data acquisition. M.D.W. participated in comparative SPR experiments and data acquisition.

