## Supplementary material for "Digital molecular kinetics enabled by large-scale nanopore arrays for multidimensional biophysical profiling": SI merterial

**Supplementary Table 1. Macroscopic kinetic measurements based on large-scale nanopore arrays.** Kinetic parameters were determined by fitting with a 1:1 binding model. Data are presented as fitted values ± standard error. Note: Because the SpyCatcher-SpyTag interaction is covalent, only the association rate constant is applicable.

| **Ligand** | **Analyte** | $K_{on}$ **($M^{-1}·s^{-1}$)** | $K_{off}$ **($s^{-1}$)** | $K_d$ **(M)** |
|---|---|---|---|---|
| CL7 | Im7-P | 5.33 (±4.34) $x10^{4}$ | 1.59 (±0.33) $x10^{-2}$ | 3.87 (±1.16) $x10^{-7}$ |
| CL7_M | Im7-P | 7.71 (±1.34) $x10^{5}$ | 1.24 (±0.22) $x10^{-3}$ | 1.71 (±0.62) $x10^{-9}$ |
| SpyCatcher | SpyTag-P | 2.02 (±0.61) $x10^{4}$ | - | - |
| Biotin | mSA-P | 9.72 (±2.85) $x10^{5}$ | 2.21 (±0.35) $x10^{-3}$ | 2.40 (±0.44) $x10^{-9}$ |
| mSA | Biotin-P | 12.88 (±6.10) $x10^{5}$ | 2.19 (±0.91) $x10^{-3}$ | 1.78 (±0.44) $x10^{-9}$ |

**Supplementary Table 2. Kinetic parameters of interactions for different molecules.**

| Molecular interaction | Methods | $K_{on}$ ($M^{-1}\cdot s^{-1}$) | $K_{off}$ ($s^{-1}$) | $K_d$ (M) |
|---|---|---|---|---|
| CL7-Im7 | Surface plasmon resonance | $1.33 \times 10^{5}$ | $6.28 \times 10^{-2}$ | $4.27 \times 10^{-7}$ |
| CL7_M-Im7 | Surface plasmon resonance | $9.24 \times 10^{5}$ | $5.04 \times 10^{-3}$ | $5.46 \times 10^{-9}$ |
| SpyCatcher-SpyTag | SDS-PAGE and Imaging Analysis[1] | $2.0\ (\pm 0.2) \times 10^{4}$ | - | - |
| Biotin-mSA | Fluorescence Polarization Spectroscopy[2] | - | - | $2.8\ (\pm 0.5) \times 10^{-9}$ (4 °C)<br>$5.5\ (\pm 0.2) \times 10^{-9}$ (37 °C) |
| CaM-M13 | Surface plasmon resonance[3] | $6.5\ (\pm 1.6) \times 10^{5}$ | $2.4\ (\pm 0.6) \times 10^{-3}$ | $3.7\ (\pm 1.8) \times 10^{-9}$ |
| GFP nanobody-GFP | Biolayer Interferometry[4] | $8.84 \times 10^{4}$ | $1.24 \times 10^{-4}$ | $1.4 \times 10^{-9}$ |
| | Surface plasmon resonance[5] | $7.68 \times 10^{5}$ | $1.74 \times 10^{-4}$ | $2.3 \times 10^{-10}$ |

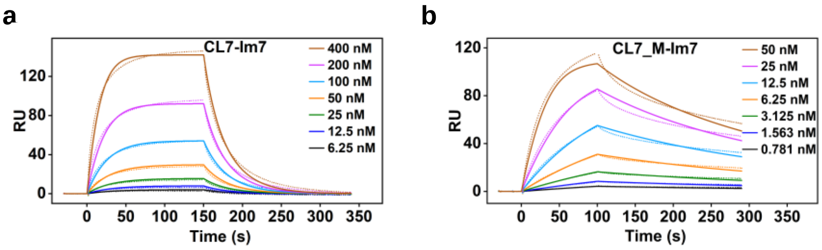


**Supplementary Fig. 1. SPR sensorgrams of CL7 or CL7_M interacting with Im7.** Sensorgrams for **(a)** CL7-Im7 and **(b)** CL7_M-Im7 display the association and dissociation phases across a series of concentrations. The dashed lines represent the experimentally measured response data, and the solid lines represent the fitted kinetic curves based on a 1:1 Langmuir binding model. These experiments were performed in 200 mM KCl and 50 mM phosphate buffer (pH 6.8).

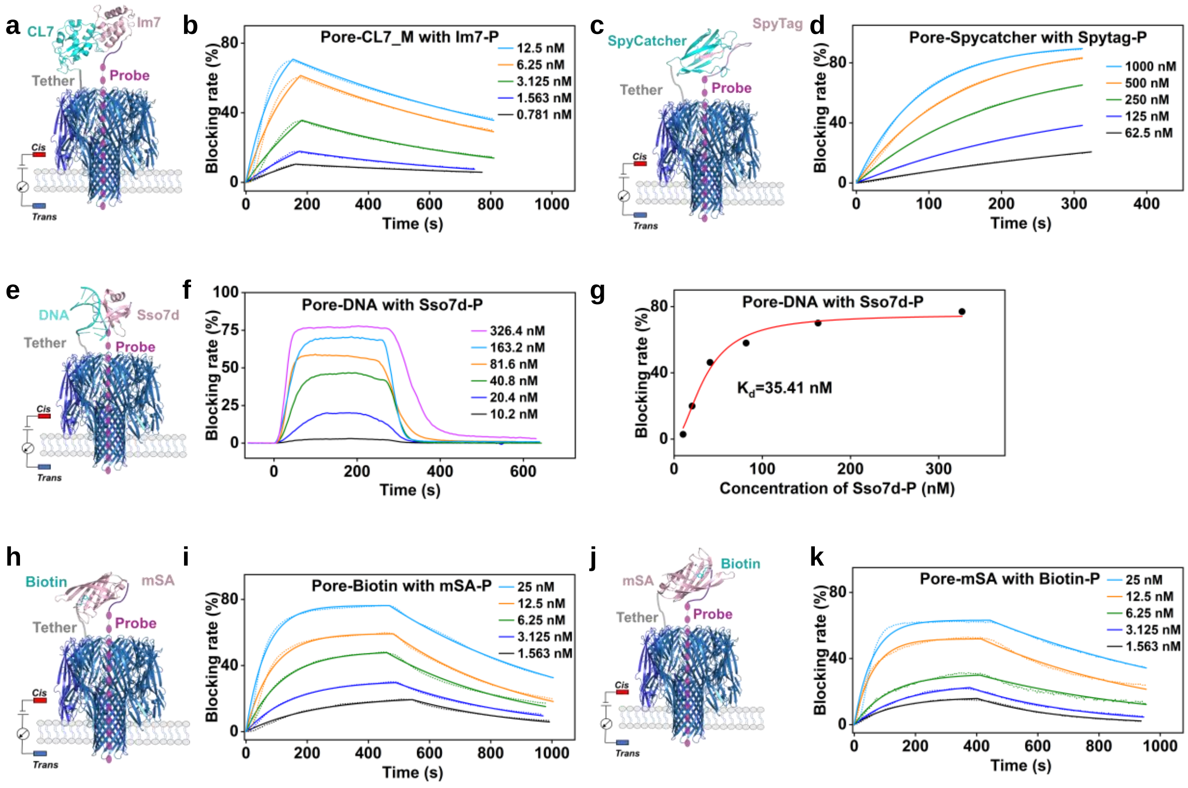


**Supplementary Fig. 2. Macroscopic kinetic measurements of diverse molecular interactions using the nanopore-probe platform. a,** Schematic illustration of the nanopore-probe strategy for detecting the CL7-Im7 interaction. **b,** Association and dissociation curves of CL7_M with Im7-P across a series of concentrations. **c,** Schematic illustration of the nanopore-probe strategy for detecting the SpyCatcher-SpyTag interaction. **d,** Association curves of SpyCatcher with SpyTag-P across a series of concentrations. Because this molecular pair undergoes covalent binding, only the association phase is shown. **e,** Schematic illustration of the nanopore-probe strategy for detecting the DNA-Sso7d interaction. **f,** Association and dissociation curves of DNA with Sso7d-P across a series of concentrations. **g,** Steady-state affinity analysis of the DNA-Sso7d interaction. The response values at various concentrations were fitted using the Hill equation to determine the $K_d$. **h,** Schematic illustration of the nanopore-probe strategy for detecting the Biotin-mSA interaction. **i,** Association and dissociation curves of Biotin with mSA-P across a series of concentrations. **j,** Schematic illustration of the nanopore-probe strategy for detecting the mSA-Biotin interaction with swapped positions of the small molecule and protein. **k,** Association and dissociation curves of mSA with Biotin-P across a series of concentrations. In the sensorgrams, dashed lines represent the experimentally measured response data (blocking rate), and solid lines represent the fitted kinetic curves based on a 1:1 binding model. These experiments were performed in 200 mM KCl, 50 mM phosphate buffer (pH 6.8) at a voltage of ±100 mV.

**Supplementary Table 3. Kinetic parameters for DNA-Sso7d, Biotin-mSA and CL7_M-Im7 interactions determined by DMK.** Corresponding kinetic parameters were calculated based on the proportions of State 1 and State 2 derived from bi-exponential fitting of the dwell time for CL7_M-Im7.

| Ligand | Analyte | Ratio of State | $K_{on}$ ($M^{-1}\cdot s^{-1}$) | $K_{off}$ ($s^{-1}$) | $K_d$ (M) |
|---|---|---|---|---|---|
| DNA | Sso7d-P | - | $7.77\ x10^{6}$ | 0.5 | $6.38\ x10^{-8}$ |
| Biotin | mSA-P | - | $3.59\ x10^{6}$ | $8.07\ x10^{-3}$ | $2.25\ x10^{-9}$ |
| CL7_M | Im7-P | State 1 (54.27%) | $1.42\ x10^{6}$ | $6.80\ x10^{-2}$ | $4.78\ x10^{-8}$ |
| | | State 2 (45.73%) | $1.20\ x10^{6}$ | $1.77\ x10^{-3}$ | $1.48\ x10^{-9}$ |

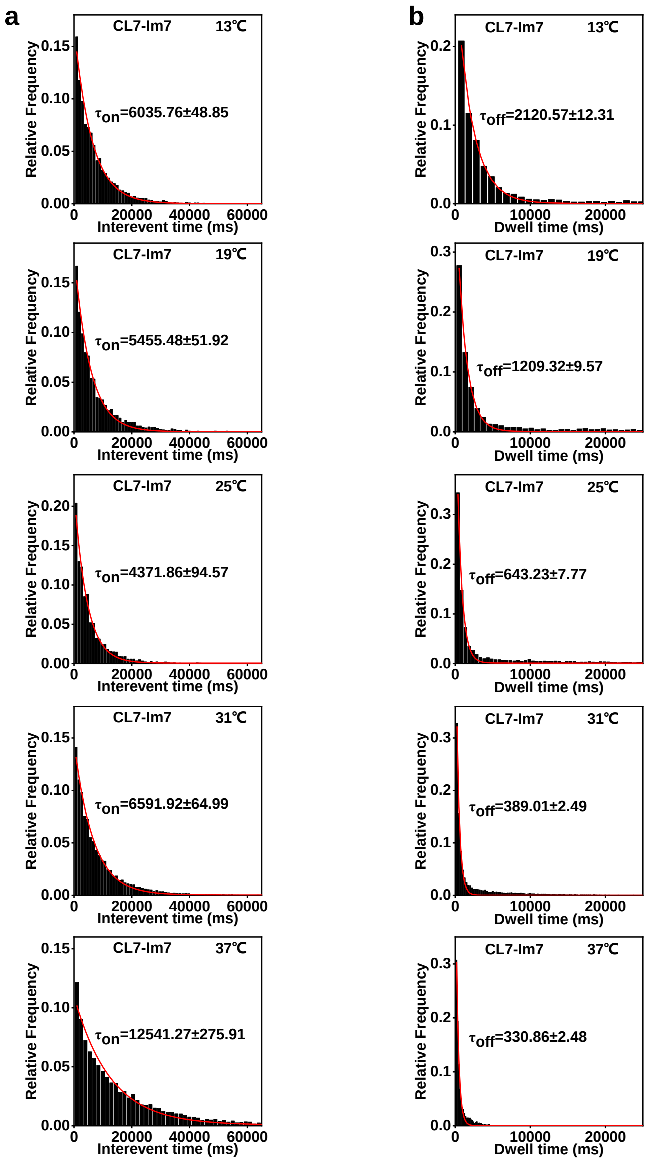


**Supplementary Fig. 3. Kinetic fitting of CL7-Im7 under different temperatures.** **a,** Statistical distributions and fitted curves of interevent time for CL7-Im7 under different temperatures. **b,** Statistical distributions and fitted curves of dwell time for CL7-Im7 under different temperatures. Both $\tau_{on}$ and $\tau_{off}$ were derived from mono-exponential fitting. These experiments were performed in 200 mM KCl, 50 mM Phosphate buffer, pH 6.8 and the voltage was ±100 mV.

**a**

$$\ln K_{\mathrm{eq}} = a + \frac{b}{T} + \frac{c}{T^2}$$

Second order polynomial (quadratic)
Y=A + B*X +C*X^2

$$\Delta H = -R\left(b + \frac{2c}{T}\right)$$

$$\Delta S = R\left(a - \frac{c}{T^2}\right)$$

**b**

| Parameters | Results |
|---|---|
| A | -260.92 |
| B | 154174.92 |
| C | -2.15E+07 |
| $R^2$ | 0.9955 |

**Supplementary Fig. 4. Fitted parameters based on the van 't Hoff equation. a,** Fitting to a second-order polynomial equation, including the calculation of thermodynamic parameters $\Delta H$ and $\Delta S$. **b,** Polynomial fitting (second-order) of the equilibrium association constants ($K_A$) derived from CL7-Im7 kinetics at different temperatures against the reciprocal of temperature, yielding fitted parameter values. Thermodynamic parameters were subsequently calculated from these fitted values using the corresponding equations.

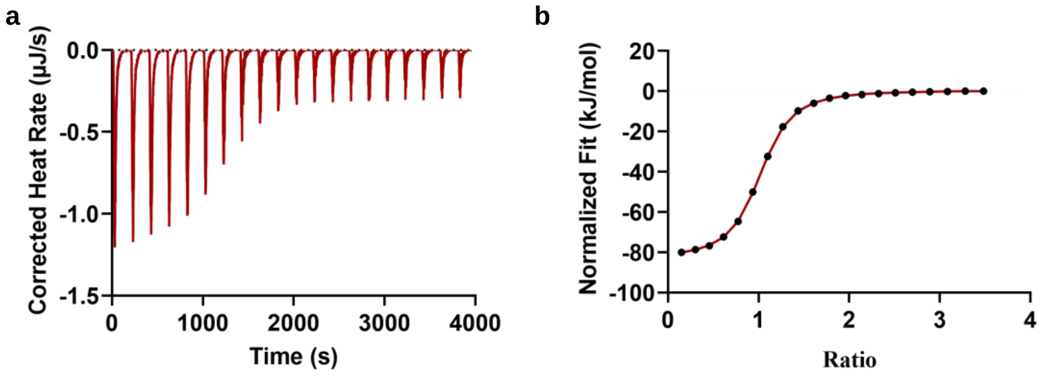


**Supplementary Fig. 5. Thermodynamic characterization of the CL7-Im7 interaction measured by ITC. a,** Raw ITC thermogram of Im7 titrated into CL7 at 25 °C. **b,** Integrated binding isotherm fitted with a one-site binding model to yield the thermodynamic parameters. These experiments were performed in 200 mM KCl, 50 mM Phosphate buffer, pH 6.8.

**Supplementary Table 4. Temperature-dependent kinetic reversibility of the CL7-Im7 interaction.** Comparison of binding kinetics measured directly at 25 °C versus those measured upon returning to 25 °C following data collection at 37 °C.

| **Ligand-Analyte** | **Temperature** | **$K_{on}$ ($M^{-1}\cdot s^{-1}$)** | **$K_{off}$ ($s^{-1}$)** | **$K_d$ (M)** |
|---|---|---|---|---|
| CL7-Im7 | 25℃ | 2.29 x$10^{6}$ | 1.55 | 6.80 x$10^{-7}$ |
| | 37-25℃ | 2.16 x$10^{6}$ | 1.42 | 6.55 x$10^{-7}$ |

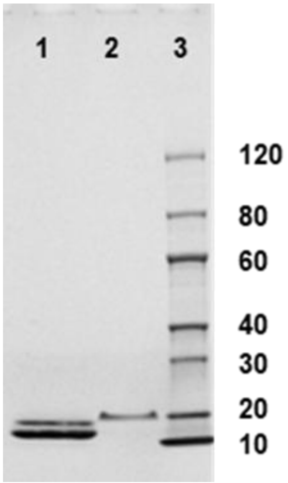


**Supplementary Fig. 6. SDS-polyacrylamide gel electrophoresis (SDS-PAGE) verifying the heterogeneity of Bim binder.** Lane 1: Bim binder. Lane 2: Bim with probe. Lane 3: Protein marker.

**Supplementary Table 5. Subpopulation fractions of heterogeneous Bim binder determined by SDS-PAGE and DMK.** Comparison of the relative proportions of Bim binder variants determined by band intensity analysis of SDS-PAGE versus channel classification based on average dwell time.

| **Type** | **Ratio** | **Channel Type** | **Ratio** |
|---|---|---|---|
| Bim binder1 | 78.49% | Short events | 73.43% |
| Bim binder2 | 21.51% | Long events | 26.57% |

**Supplementary Table 6. Single-molecule kinetic analysis of heterogeneous Bim binder interacting with Bim by DMK.** Kinetic parameters ($K_{on}$, $K_{off}$, and $K_d$) for Bim binder 1-Bim and Bim binder 2-Bim subpopulations (extrapolated to V=0), derived from single-exponential fits of interevent time and dwell time distributions following channel categorization based on the average dwell time threshold.

| **[Bim binder-Bim]** | $K_{on}$ **($M^{-1}\cdot s^{-1}$)** | $K_{off}$ **($s^{-1}$)** | $K_d$ **(M)** |
|---|---|---|---|
| [Bim binder 1 -Bim] | $1.00 \times 10^{6}$ | $4.14 \times 10^{-2}$ | $4.13 \times 10^{-8}$ |
| [Bim binder 2 -Bim] | $1.92 \times 10^{6}$ | $3.96 \times 10^{-3}$ | $2.07 \times 10^{-9}$ |

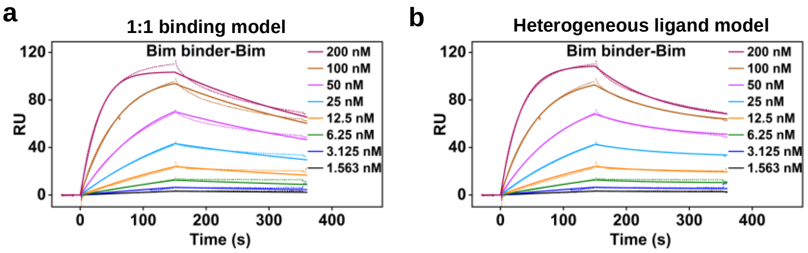


**Supplementary Fig. 7. SPR sensorgrams and kinetic fitting of the Bim binder-Bim interaction under different binding models. a,** Kinetic fitting of concentration-series sensorgrams using a 1:1 Langmuir binding model. **b,** Kinetic fitting of concentration-series sensorgrams using a heterogeneous ligand model. Dashed lines represent experimental SPR sensorgrams at indicated Bim concentrations (1.563–200 nM), and colored solid lines represent the corresponding kinetic fits.

**Supplementary Table 7. Kinetic parameters of the Bim binder-Bim interaction determined by surface plasmon resonance (SPR) using different binding models.** Summary of fitted association rate constants ($K_{on}$), dissociation rate constants ($K_{off}$), and equilibrium dissociation constants ($K_d$) derived from 1:1 binding and heterogeneous ligand models.

| Model | $K_{on1}$ ($M^{-1}\cdot s^{-1}$) | $K_{off1}$ ($s^{-1}$) | $K_{d1}$ (M) | $K_{on2}$ ($M^{-1}\cdot s^{-1}$) | $K_{off2}$ ($s^{-1}$) | $K_{d2}$ (M) | State1:State2 |
|---|---|---|---|---|---|---|---|
| 1:1 binding | $2.22 \times 10^{5}$ | $2.45 \times 10^{-3}$ | $1.10 \times 10^{-8}$ | - | - | - | - |
| Heterogeneous ligand | $2.22 \times 10^{5}$ | $6.04 \times 10^{-7}$ | $2.72 \times 10^{-12}$ | $1.13 \times 10^{5}$ | $9.76 \times 10^{-3}$ | $8.64 \times 10^{-8}$ | 47.5% : 52.5% |

**Supplementary Table 8. Extrapolated zero-voltage kinetic parameters of CL7-Im7 and CaM-M13 interactions determined by DMK.** Kinetic parameters for State 1 and State 2 were derived from double-exponential fitting of dwell time distributions across an applied voltage gradient. For CL7-Im7, zero-voltage parameters represent mean values across all voltages due to the absence of voltage dependence. For CaM-M13, zero-voltage $K_{on}$ represents the mean across voltages, whereas zero-voltage $K_{off}$ and $K_d$ were extrapolated via linear regression to the y-intercept (Equation 2).

| Molecular interaction | Ratio of State (100 mV) | $K_{on}$ ($M^{-1}\cdot s^{-1}$) | $K_{off}$ ($s^{-1}$) | $K_d$ (M) |
|---|---|---|---|---|
| [CL7-Im7] | State 1 (64.65%) | $1.31 \times 10^{6}$ | 1.53 | $1.38 \times 10^{-6}$ |
| | State 2 (35.35%) | $8.23 \times 10^{5}$ | $5.21 \times 10^{-2}$ | $7.35 \times 10^{-8}$ |
| [CaM-M13] | State 1 (28.21%) | $1.78 \times 10^{6}$ | $1.26 \times 10^{-1}$ | $7.08 \times 10^{-8}$ |
| | State 2 (71.79%) | $3.80 \times 10^{6}$ | $4.75 \times 10^{-3}$ | $1.25 \times 10^{-9}$ |

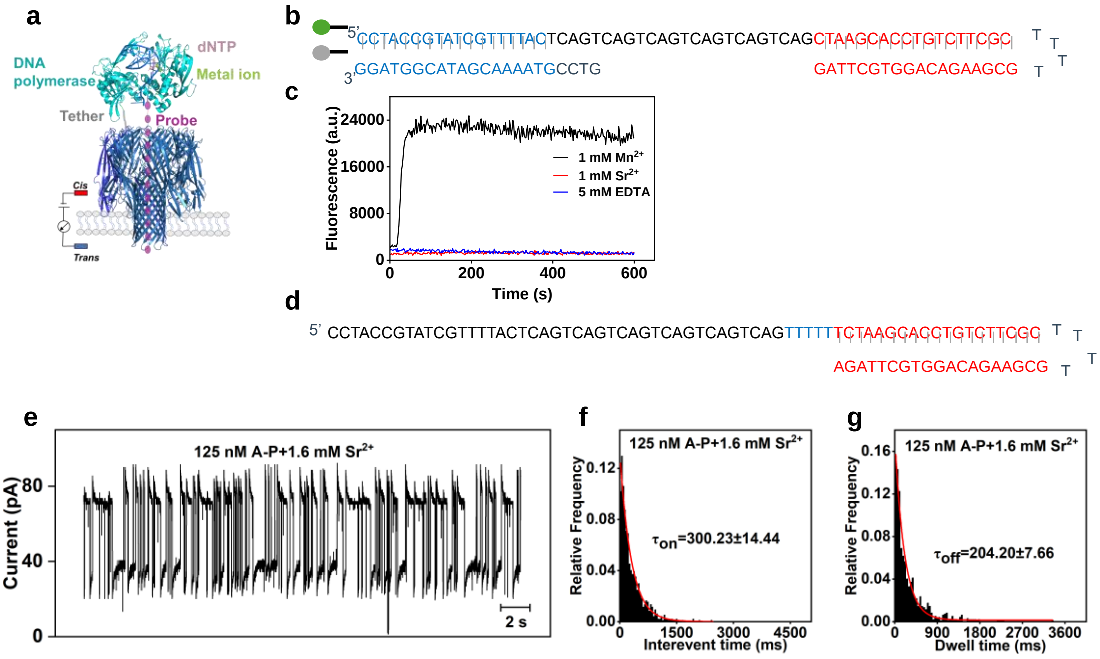


**Supplementary Figure 8. DMK-based kinetic analysis of transient interactions between the elongation complex and complementary nucleotides. a,** Schematic of the nanopore-tethered strategy for detecting interactions between the elongation complex and complementary nucleotides. **b-c,** Fluorometric assay for DNA polymerase activity. **b,** Schematic of the activity assay using a hairpin nucleic acid substrate labeled with a 5-prime fluorophore (Cy3) and a 3-prime quencher (BHQ2). **c,** Fluorescence responses of DNA polymerase in the presence of 1 mM various metal ions (MnCl2 and SrCl2) or 5 mM EDTA. **d,** Sequence structure of the hairpin nucleic acid used for DNA polymerase binding, featuring a starting binding site of 5 thymine residues at the 3-prime end. **e-g,** Single-molecule kinetic characterization under 1.6 mM Sr2+ conditions with 125 nM dA6P-P. **e,** Representative ionic current trace of elongation complex binding events. f,g, Statistical distributions and single-exponential fits for interevent time **(f)** and dwell time **(g)**. Zero-voltage kinetic parameters ($K_{on}$ = 2.66 x $10^7$ $M^{-1}$. $s^{-1}$, $K_{off}$ = 4.90 $s^{-1}$, and $K_d$ = 183.78 nM) were calculated from the derived $\tau_{on}$ and $\tau_{off}$ values. All experiments were performed in buffer containing 650 mM KAc and 50 mM Phosphate buffer (pH 6.8) at an applied voltage of +70 mV.

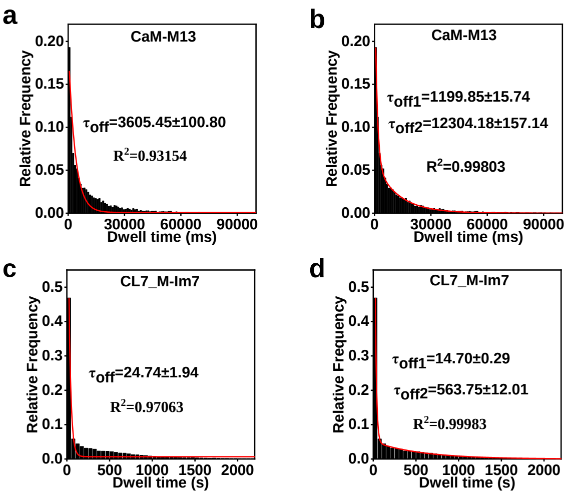


**Supplementary Figure 9. Comparison between single-exponential and double-exponential fits for residence time distributions. a,b,** Statistical dwell time distributions and corresponding single-exponential fit (**a,** R2 = 0.932) and double-exponential fit (**b,** R2 = 0.998) for CaM-M13 interactions. **c,d,** Statistical dwell time distributions and corresponding single-exponential fit (**c,** R2 = 0.971) and double-exponential fit (**d,** R2 = 0.999) for CL7_M-Im7 interactions. These experiments were performed in 200 mM KCl, 50 mM Phosphate buffer, pH 6.8 and the voltage was ±100 mV.

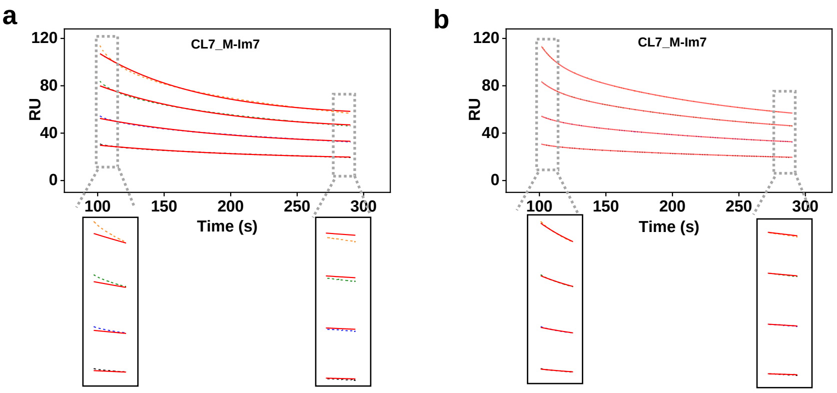


c

| [CL7_M-Im7] (nM) | $R^2$ (Single Exponential Model) | $R^2$ (Double Exponential Model) |
|---|---|---|
| 50 | 0.99305 | 0.99992 |
| 25 | 0.99526 | 0.99993 |
| 12.5 | 0.99627 | 0.99991 |
| 6.25 | 0.99652 | 0.99969 |

**Supplementary Figure 10. Comparison of single-exponential and double-exponential fitting for the dissociation curves of the CL7_M-Im7 measured by SPR. a,** Sensorgram of the dissociation phase from the SPR measurement, with the single-exponential model used for fitting. **b,** Sensorgram of the dissociation phase from the SPR measurement, with the double-exponential model used for fitting. The dashed lines represent the experimentally measured response data, and the solid lines represent the fitted kinetic curves. Enlarged views of the corresponding sensorgrams are shown below each panel. **c,** Comparison of $R^2$ between single-exponential and double-exponential fitting.
